# Multiscale structural MRI differences among collegiate American football players, cross-country runners, and non-athletes

**DOI:** 10.64898/2026.10.01.756138

**Authors:** Arunim Guchait, Chiao-Yun Chen, Neil G. Muggleton

## Abstract

Structural differences observed in collision-sport athletes may reflect repetitive head-impact exposure, broader effects of athletic training, or pre-existing individual variation, making inclusion of a non-collision athlete group important for interpretation of any differences seen. Here, structural T1-weighted MRI data from 17 collegiate American football players, 16 cross-country runners, and 9 non-athletes were analyzed using FSL voxel-based morphometry (VBM), FIRST subcortical volumetry and vertex-wise shape analysis, and FAST whole-brain tissue segmentation. Spatial inference used permutation testing with threshold-free cluster enhancement and family-wise error correction, while scalar analyses used multiplicity-corrected procedures. FAST-derived absolute tissue volumes were also examined in a sensitivity analysis adjusting for FreeSurfer estimated total intracranial volume (eTIV). VBM showed recurrent brainstem-centered differences, with non-athletes showing greater local gray matter measures than both athlete groups, whereas football players showed greater brainstem measures than runners. FIRST vertex-wise analysis identified corrected differences involving the brainstem and selected subcortical structures, while no total FIRST structure volume survived false discovery rate correction. Unadjusted FAST analysis identified a CSF group effect, *F* (2, 24.38) = 11.46, *p* < .001, *q* = .001. After eTIV adjustment, the CSF group effect remained, *F* (2, 38) = 14.30, *q* < .001, and a white matter group effect was also identified, *F* (2, 38) = 9.59, *q* < .001; gray matter and total segmented tissue did not show corrected adjusted effects. These findings indicate that structural variation depends on anatomical scale, comparison group, and head-size adjustment and cannot be attributed uniquely to collision-sport exposure.

## 1 Introduction

Participation in American football provides a particularly important context in which to examine the relationship between long-term sporting exposure and brain structure. Unlike many forms of physical activity, football involves repeated acceleration and deceleration of the head during routine practices and competition, arising from both direct head contact and forces transmitted through impacts elsewhere on the body. Of relevance here is the distinction between clinically diagnosed concussion and repetitive head impacts that do not produce recognizable acute symptoms. The latter are commonly described as subconcussive or subclinical head impacts and may occur repeatedly without meeting diagnostic criteria for concussion (Koerte et al., 2023). The absence of an acute clinical diagnosis does not necessarily imply an absence of biomechanical exposure or measurable neural change. Prospective studies have reported MRI differences across a football season in athletes who did not sustain a clinically evident concussion, including white matter changes associated with helmet-derived impact measures and alterations detected across multimodal neuroimaging measures (Bazarian et al., 2014; Slobounov et al., 2017). These observations do not establish that routine football participation produces lasting brain injury, but they provide a rationale for examining whether structural differences are detectable in athletes with prolonged exposure to the sport.

A difficulty with this question, however, is that American football players differ from non-athletes in many respects other than exposure to head impacts. Competitive athletes typically experience sustained physical training, repeated motor learning, structured practice, high training loads, and distinctive physiological and behavioral demands. Some of these characteristics may themselves be associated with brain structure differences. Neuroimaging studies of endurance athletes, for example, have identified structural and functional differences between trained runners and non-running controls, illustrating that measurable brain differences can occur in sporting populations without routine collision exposure (Cao et al., 2021). This means that a structural difference between football players and non-athletes cannot automatically be attributed to repetitive head impacts. The same observation could reflect physical fitness, long-term training, sport-related skill acquisition, developmental or lifestyle differences, pre-existing characteristics that influence selection into sport, or some combination of these factors. In other words, the conventional football-versus-non-athlete comparison combines collision-sport exposure with the broader distinction between being a highly trained collegiate athlete and not being one. A more informative design therefore requires an athlete group that shares substantial sporting engagement but does not share the same routine collision profile.

Cross-country runners provide such a comparison and consequently occupy a central rather than secondary role in the study here. The source dataset includes American football players, cross-country runners, and non-athletes, allowing three related questions to be considered within the same cohort (Caron et al., 2021). The football-versus-non-athlete comparison tests whether the collision-sport group differs from individuals without collegiate athletic participation, but any difference in this contrast contains both athletic-status and football-related components. The runner-versus-non-athlete comparison provides a parallel test of whether structural differences are also present in athletes whose training is intensive but does not involve routine collision exposure. Most importantly, the football-versus-runner comparison reduces the broad athlete-versus-non-athlete distinction and provides a more informative contrast for identifying patterns that may be more specific to American football. It is important to note, though, that this logic does not completely isolate the effect of repetitive head impacts because football and endurance running differ in many additional physiological and behavioral respects. Nevertheless, the three-group structure provides a stronger interpretive framework than a football-versus-control comparison alone. Evidence that is shared by both athlete groups would be more consistent with general athletic or training-related factors (or other characteristic differences related to voluntary engagement in significant sport or exercise engagement), whereas differences between the two athlete groups would motivate consideration of sport-specific exposures, one aspect of which relates to effects of collisions.

A second issue is that structural variation may not be expressed at only one anatomical scale. For this reason, reliance on a single morphometric measure could provide an incomplete account of group differences. Voxel-based morphometry (VBM) enables whole-brain voxel-wise assessment of local gray matter variation after tissue segmentation and spatial normalization and is therefore useful when regional differences are not restricted to predefined anatomical structures (Ashburner & Friston, 2000). FIRST addresses a different question by using a Bayesian model of shape and appearance to segment subcortical structures (Patenaude et al., 2011). Its volumetric measures quantify the overall size of individual structures, whereas vertex-wise analysis can identify localized surface displacement that may be present even when total structure volume is similar between groups. FAST provides a broader level of description by segmenting structural MRI into tissue classes, including cerebrospinal fluid, gray matter, and white matter, using probabilistic tissue classification (Zhang et al., 2001). These approaches therefore address related but non-equivalent properties of the same T1-weighted images. Agreement across methods can indicate that an anatomical pattern is detectable at more than one measurement scale, whereas disagreement can itself be informative by showing that a group difference is localized, shape-dependent, or apparent only at the level of global tissue composition.

The study here uses the publicly available collegiate athlete dataset described by Caron et al. (2021), which was released as an open and reproducible neuroimaging resource containing two collegiate athlete groups and a non-athlete group and emphasizing diffusion MRI derivatives, tractography, and structural connectivity. The available T1-weighted anatomical images make it possible to address a complementary set of structural-morphometric questions that were not the primary focus of the original data descriptor. Three aims were therefore addressed. The first was to determine whether localized differences in modulated gray matter volume were detectable among American football players, cross-country runners, and non-athletes using FSL-VBM. The second was to determine whether subcortical structures differed in either total volume or localized surface morphology using FIRST. The third was to determine whether the groups differed in global cerebrospinal fluid, gray matter, or white matter volume using FAST. Together, these analyses were intended to characterize structural variation across voxel-wise, structure-specific, and whole-brain tissue scales rather than to assume that any football-associated effect would appear in a particular region or direction. It was expected that comparison with both non-athletes and non-collision athletes would help distinguish patterns associated with collegiate athletic participation more generally from those that appeared relatively more specific to the American football group. Given the cross-sectional design and the absence of complete lifetime exposure information, however, any such differences were treated as anatomical associations rather than evidence that football participation caused structural change.

## 2 Materials and Methods

### 2.1 Dataset, participants, and ethics

This study performed a secondary analysis using open-access structural magnetic resonance imaging data derived from a collegiate athlete dataset. Originally acquired at Indiana University, the dataset was subsequently made available via brainlife.io (Caron et al., 2021). The source dataset contains anatomical T1-weighted and diffusion MRI data from collegiate American football players, cross-country runners, and non-athlete university students and was developed to support open and reproducible investigation of long-term athletic exposure and the brain. The data used for the present analysis are publicly available through the brainlife.io repository under DOI 10.25663/brainlife.pub.14.

The released sample comprised 42 male participants: 17 American football players, 16 cross-country runners, and 9 non-athletes. All eligible T1-weighted datasets were retained for the structural analyses reported here. The American football group consisted of fourth- and fifth-year varsity Indiana University football starters and had a reported mean age of 21.1 years (SD = 1.5). Twenty-one football players originally entered the study; one did not complete the study and three did not complete the diffusion MRI acquisition, leaving 17 participants in the released dataset. Potential football participants were excluded from the original study if they reported a diagnosed concussion within six months preceding enrollment. However, three of the retained football players had a more remote history of diagnosed concussion, with two having sustained a concussion approximately three years before data collection and one approximately two years before data collection. Importantly, information concerning concussion history before entry to Indiana University was not collected, meaning that lifetime exposure to diagnosed concussion could not be established.

The cross-country group initially included 19 members of the Indiana University cross-country team, with a reported mean age of 20.2 years (SD = 2.5). These participants were included as a non-collision athlete group and were matched to the football players on age and collegiate experience. Three runners had unusable anatomical T1-weighted images, leaving 16 participants. The non-athlete group was drawn from an undergraduate Psychology and Neuroscience subject pool and had a reported mean age of 19.9 years (SD = 3.0). Eleven individuals were initially selected on the basis of age, sex, and estimated family income; two were excluded from the released dataset because their diffusion data contained artifacts that could not be corrected, leaving nine non-athletes. The source study did not collect additional information concerning habitual exercise in this group.

The original data collection was approved by the Indiana University Institutional Review Board, and all participants provided written informed consent. Because the analyses reported here were conducted retrospectively using publicly available, de-identified neuroimaging data, no new participant recruitment or data acquisition was undertaken for the present study.

### 2.2 MRI acquisition

MRI data were acquired at the Indiana University Imaging Research Facility using a 3-Tesla Siemens TIM Trio scanner equipped with a 12-channel head coil. The structural analyses reported here used the T1-weighted anatomical image acquired for each participant. The anatomical sequence had a repetition time of 1800 ms, echo time of 2.67 ms, inversion time of 900 ms, flip angle of 9°, and receiver bandwidth of 150 Hz/pixel. Images comprised 160 sagittal slices with a 256-mm field of view and a 256 × 256 acquisition matrix. Slice thickness was 1 mm, resulting in an isotropic spatial resolution of 1 × 1 × 1 mm. Only the T1-weighted anatomical images were used for the morphometric analyses described below; the diffusion and functional MRI acquisitions contained within the larger source dataset were not analyzed in the present study.

### 2.3 Structural-analysis overview

The T1-weighted anatomical images were analyzed using three complementary structural MRI pipelines implemented within the FMRIB Software Library (FSL; Jenkinson et al. (2012); Smith et al. (2004)). First, voxel-based morphometry (VBM) was used to assess voxel-wise differences in modulated gray matter volume. Second, subcortical segmentation and shape analysis were performed using FIRST to derive both volumetric and vertex-wise measures of deep gray matter structures. Third, FAST tissue segmentation was used to estimate global cerebrospinal fluid (CSF), gray matter, and white matter volumes using partial-volume modeling. Together, these approaches provided converging evidence across voxel-wise, structure-specific, and whole-brain tissue scales.

### 2.4 FSL voxel-based morphometry

Whole-brain gray matter morphology was examined using the FSL-VBM pipeline, an optimized voxel-based morphometry framework implemented within FSL (Ashburner & Friston, 2000; Douaud et al., 2007; Smith et al., 2004). VBM allows regional gray matter differences to be tested on a voxel-wise basis following tissue segmentation, nonlinear registration, modulation, and spatial smoothing.

Initial preprocessing involved brain extraction of each T1-weighted image using the FSL-VBM brain-extraction stage, with outputs visually inspected to ensure adequate removal of non-brain tissue. Because the three participant groups differed in sample size, the VBM analyses were organized as five balanced pairwise comparisons, with a separate study-specific gray matter template generated for each comparison. This ensured equal representation of the two groups contributing to each template, consistent with FSL-VBM guidance for reducing group-related registration bias during study-specific template construction.

For the American football player-versus-non-athlete analyses, the nine non-athletes were compared separately with the first nine American football players and the last nine American football players according to participant ordering in the source dataset. Each comparison therefore used a study-specific template constructed from nine American football players and nine non-athletes (*n* = 18). Similarly, the nine non-athletes were compared separately with the first nine and last nine cross-country runners, with each corresponding template constructed from nine runners and nine non-athletes (*n* = 18). The athlete groups contained fewer than 18 participants and the two football subsets were not completely independent, with one football player included in both subsets, and the two runner subsets overlapped by two participants. Thus, for the direct athlete comparison, 16 American football players were compared with all 16 cross-country runners, and the corresponding study-specific template was constructed from those 32 participants. This meant each statistical comparison used a template containing equal representation of the two groups in that analysis.

For each analysis, the gray matter images defined in the corresponding analysis-specific template construction list (template_list) were registered to standard space and averaged. The averaged image was then left-right reflected and re-averaged to produce a symmetric study-specific gray matter template. The gray matter images entering each comparison were subsequently nonlinearly registered to the corresponding study-specific template. To preserve local volumetric information following spatial normalization, the registered images were modulated using the Jacobian determinants of the deformation fields, such that voxel intensities reflected local tissue volume rather than tissue concentration alone.

The modulated gray matter images were concatenated into a four-dimensional dataset and spatially smoothed within the FSL-VBM framework. Final statistical inference used the smoothed modulated gray matter dataset with a 3 mm Gaussian kernel (*σ* = 3 mm; *GM_mod_merg_s3*). The smoothing configuration was selected before final hypothesis testing on the basis of the predefined processing workflow and inspection of the spatial stability of the resulting maps. The same smoothing specification was applied consistently across the five VBM analyses.

Voxel-wise statistical inference was performed separately for each balanced comparison using FSL’s permutation-based nonparametric inference tool (randomise) with 5,000 permutations and threshold-free cluster enhancement (TFCE), which avoids the need to specify an arbitrary initial cluster-forming threshold (Smith & Nichols, 2009). Two directional contrasts were specified for each pairwise analysis. Corrected statistical maps were evaluated at a family-wise error-corrected threshold of *p ≤* .05.

### 2.5 FIRST segmentation, volumetry, and vertex-wise shape analysis

Subcortical segmentation and shape analysis were performed using FSL FIRST, which applies a Bayesian shape and appearance model to segment deep gray matter structures from T1-weighted images (Patenaude et al., 2011). Fifteen structures were included: bilateral nucleus accumbens, amygdala, caudate, hippocampus, pallidum, putamen, thalamus, and the brainstem.

For each participant, subcortical structures were automatically segmented using the standard FIRST processing pipeline (run_first_all), and all outputs were visually inspected for anatomical plausibility and registration accuracy. Quality control was performed prior to any quantitative analysis, and representative segmentations are provided in Supplementary Figure S1.

Structure-wise volumes were extracted directly from the segmentation outputs and used for group-level comparisons across all 15 regions. These volumetric analyses provided global measures of structural differences but did not capture spatially localized shape variation.

To address this limitation, vertex-wise shape analysis was performed using the FIRST shape model. Individual shape parameters (.bvars files) were concatenated across participants in a fixed group order, and group-level design matrices were constructed for each comparison. Surface reconstructions were generated in MNI space, and vertex-wise displacement maps were derived using FIRST’s shape projection framework (implemented via first_utils). This procedure yields signed surface displacement values along vertex normals, enabling localized shape differences to be statistically evaluated.

Permutation-based inference was performed using 10,000 permutations with TFCE correction. Both directional t-contrasts and omnibus F-contrasts were evaluated for each structure and group comparison. Corrected statistical maps were thresholded at FWE p *≤* .05.

### 2.6 FAST whole-brain tissue segmentation

Whole-brain tissue volumes were estimated using FSL FAST, which performs probabilistic segmentation using a hidden Markov random field model with expectation-maximization optimization (Zhang et al., 2001). Prior to segmentation, images were skull-stripped and bias-field corrected to improve tissue classification accuracy.

FAST was configured for three-class segmentation (CSF, gray matter, white matter) with partial-volume estimation. This approach allows each voxel to contain fractional tissue contributions, improving volumetric accuracy in boundary regions. Tissue volumes were computed by integrating partial-volume estimates across all voxels and multiplying by voxel dimensions. Total segmented tissue volume was defined as the sum of CSF, gray matter, and white matter volumes. Because the FAST outcomes represented absolute tissue volumes, a separate sensitivity analysis was performed to determine whether the observed group effects were retained after adjustment for FreeSurfer estimated total intracranial volume (eTIV) obtained from the source-dataset metadata. FreeSurfer eTIV is derived from atlas-based scaling and provides an automated estimate of intracranial size (Buckner et al., 2004).

### 2.7 Statistical analysis and multiple-comparison control

Voxel-wise analyses (VBM and FIRST shape) were performed using nonparametric permutation testing with TFCE correction, implemented in FSL randomise. VBM analyses used 5,000 permutations, while FIRST shape analyses used 10,000 permutations. Family-wise error correction was applied at p *≤* .05.

Volumetric analyses derived from FIRST were evaluated using Welch’s t-tests, with false discovery rate (FDR) correction applied across the 15 structures within each group comparison. Effect sizes are reported as Hedges’ g with 95% confidence intervals.

FAST-derived absolute tissue volumes were evaluated initially using three-group Welch analyses of variance followed by pairwise Welch’s *t* -tests. Benjamini–Hochberg FDR correction was applied across the four omnibus FAST tissue-volume tests and separately across the 12 pairwise tissue-volume comparisons. Pairwise effect sizes were reported as Hedges’ *g*with 95% confidence intervals. Across the scalar analyses, inference was based on multiplicity-corrected results rather than uncorrected *p* values or effect-size magnitude alone.

A head-size sensitivity analysis was additionally performed for the four FAST-derived absolute tissue volumes using FreeSurfer estimated total intracranial volume (eTIV) from the source dataset. eTIV was converted to cm^3^ and centered at the sample mean of 1703.345 cm^3^. For each FAST outcome, an ordinary least-squares regression model included group and centered eTIV as predictors. Homogeneity of regression slopes were evaluated before the adjusted analysis by including group-by-eTIV interaction terms. To retain robustness to unequal residual variances and unequal group sizes, inferential statistics were calculated using HC3 heteroskedasticity-consistent covariance estimates (MacKinnon & White, 1985). The adjusted omnibus group effect was evaluated using a joint robust Wald *F* test, and pairwise adjusted group contrasts were estimated at the grand mean eTIV with robust 95% confidence intervals. Benjamini-Hochberg false discovery rate correction was applied across the four adjusted omnibus tests and separately across the 12 adjusted pairwise comparisons (Benjamini & Hochberg, 1995). The eTIV-adjusted models were treated as sensitivity analyses of the FAST findings.

Analysis scripts implementing the structural MRI processing and statistical procedures described above are publicly available through GitLab at https://gitlab.com/brain-behav-lab/athlete-structural-mri.

## 3 Results

### 3.1 Whole-brain voxel-based morphometry

The clearest pattern in the whole-brain VBM analysis was the repeated identification of brainstem-centered differences across the balanced group comparisons. The complete corrected cluster output is provided in Supplementary Table S1, while the principal spatial findings are illustrated in Figure 1.

**Figure 1:**
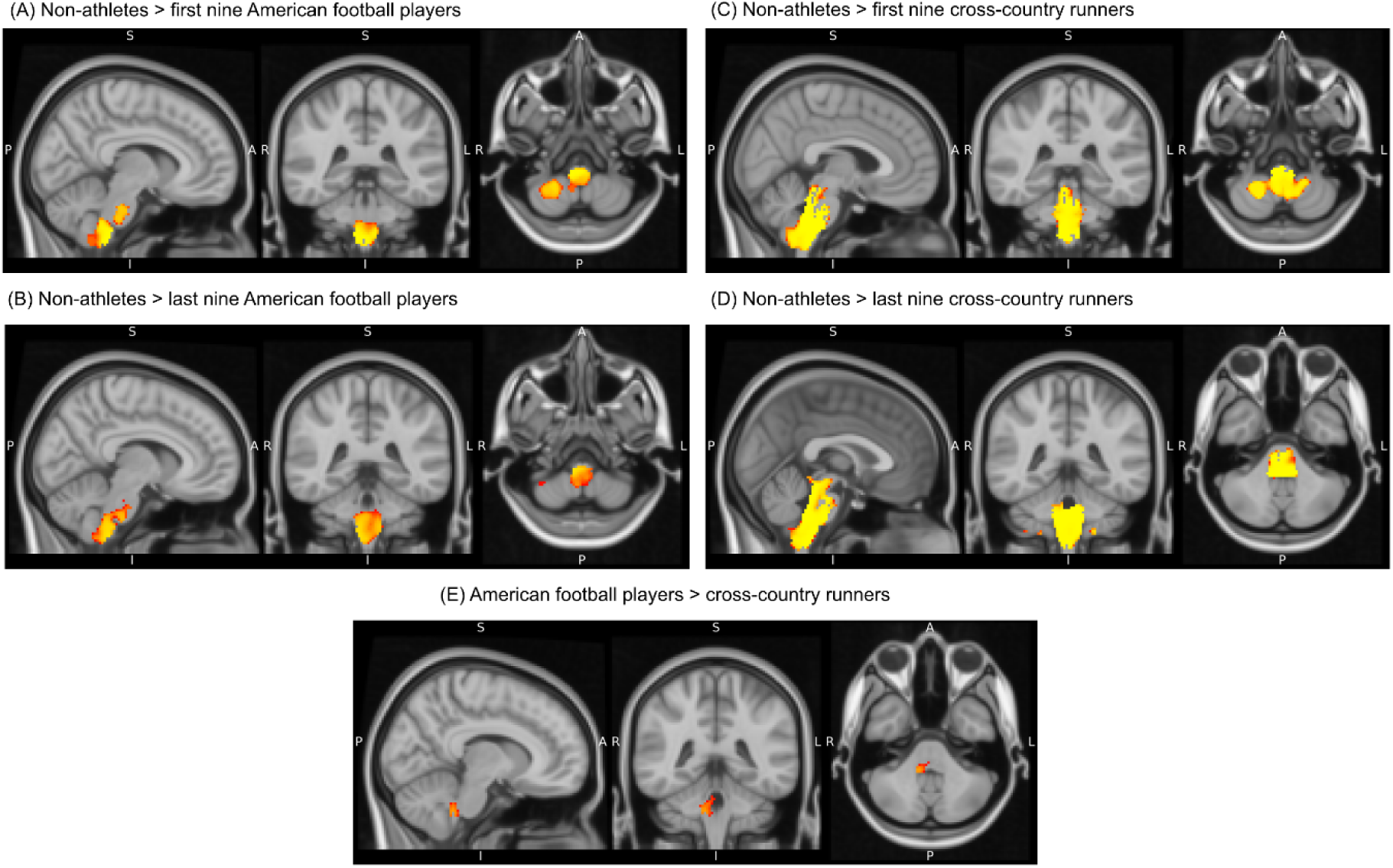
Primary whole-brain VBM results. Red-yellow overlays show significant regional differences in modulated gray matter volume from FSL-VBM analyses, displayed on the MNI152 T1 2-mm template. Maps show TFCE-corrected permutation-test results thresholded at corrected 1-p *≥* 0.95, equivalent to corrected p *≤* 0.05. Panels correspond to the following contrasts and display locations: (A) Non-athletes > first nine American football players, centered at x = 8, y = -38, z = -58; (B) Non-athletes > last nine American football players, centered at x = 8, y = -40, z = -58; (C) Non-athletes > first nine cross-country runners, centered at x = 4, y = -40, z = -62; (D) Non-athletes > last nine cross-country runners, centered at x = 0, y = -42, z = -66; and (E) American football players > cross-country runners, centered at x = 10, y = -42, z = -40. All coordinates are in MNI152 space. The display range displayed in all panels is set to 0.95-1.00 using the red-yellow color map.

In the first balanced comparison of nine American football players with the nine non-athletes, a large cluster was present in the non-athletes > American football players direction and was centered in the brainstem. This cluster contained 1,560 voxels, with a maximum corrected 1 *− p* value of 1.000, TFCE-corrected *p* < .001, peak *t* = 12.60, and peak MNI coordinates [8, *−*38, *−*58]. A second corrected cluster was identified in the right cerebral cortex, but this was limited to three voxels, TFCE-corrected *p* = .042, peak *t* = 7.37, at [28, 46, 24]. The second balanced football comparison showed the same broad brainstem difference. Non-athletes showed greater modulated gray matter volume than the last nine football players in a 1,165-voxel brainstem cluster, TFCE-corrected *p* = .005, peak *t* = 6.73, at [8, *−*40, *−*58]. An additional corrected cluster in right cerebellar lobule VIIIa contained only nine voxels was seen with TFCE-corrected *p* = .045, peak *t* = 6.32, at [38, *−*52, *−*60].

The runner comparisons produced a similarly brainstem-centered pattern. For the first nine cross-country runners versus non-athletes, the non-athletes > runners contrast identified a 3,228-voxel brainstem cluster, TFCE-corrected *p* < .001, peak *t* = 10.50, with the corrected-map peak at [4, *−*40, *−*62]. In the last-nine runner comparison, a still larger brainstem cluster of 3,964 voxels was present in the same non-athletes > runners direction, TFCE-corrected *p* < .001, peak *t* = 10.60, at [0, *−*42, *−*66].

The direct comparison between the two athlete groups also showed differences. American football players showed greater modulated gray matter volume than cross-country runners in a 275-voxel brainstem cluster, TFCE-corrected *p* = .019, peak *t* = 5.30, at [10, *−*42, *−*40]. A second brainstem cluster in the same direction contained only two voxels and lay at the corrected threshold, TFCE-corrected *p* = .050, peak *t* = 3.83. No corrected clusters were significant in the reverse direction. Thus, although the direction depended on the groups being contrasted, the spatially substantial VBM findings were consistently concentrated in the brainstem rather than being broadly distributed throughout cortical gray matter.

### 3.2 FIRST vertex-wise shape analysis

The FIRST vertex-wise analysis identified FWE-corrected local shape differences in all three pairwise group comparisons. Table 1 summarizes the corrected clusters, and Figures 2–4 show the corresponding statistical maps. In contrast to the total-volume analysis described below, these vertex-wise effects survived spatial correction for multiple comparisons.

**Figure 2:**
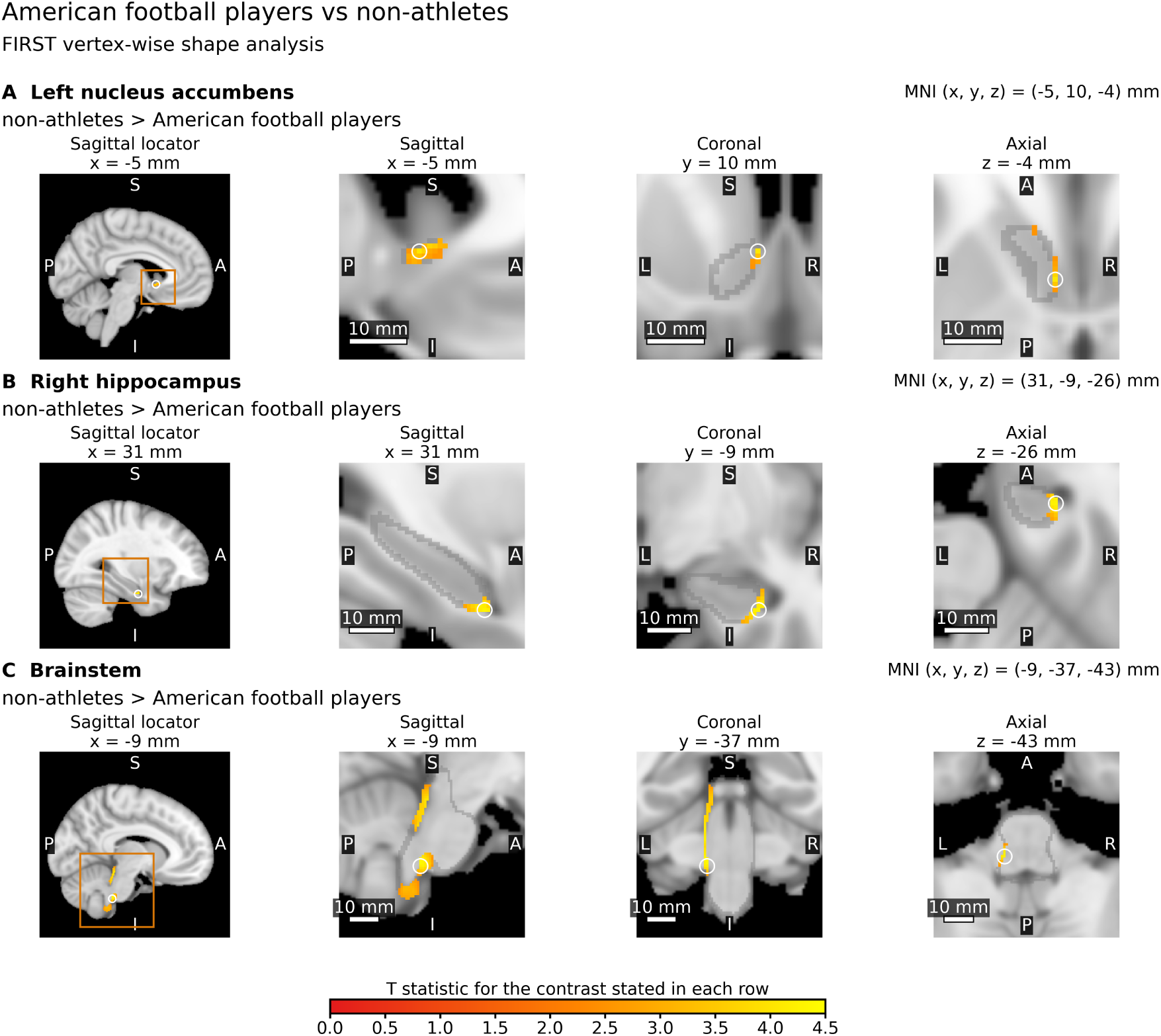
FIRST vertex-wise shape differences between American football players and non-athletes. Significant local surface differences were identified in the left nucleus accumbens (A), right hippocampus (B), and brainstem (C), all for the non-athletes > American football players contrast. Red-to-yellow overlays show directional t statistics at vertices surviving TFCE-based family-wise error correction (pFWE *≤* .05; 10,000 permutations). Each row shows a sagittal locator together with enlarged sagittal, coronal, and axial views intersecting the reported Montreal Neurological Institute (MNI) coordinate. Orange boxes in the locator images indicate the field of view of the enlarged sagittal view. Gray outlines delineate the tested FIRST structure, and white circles mark the reported MNI coordinate. Images are displayed on the MNI152 1-mm T1-weighted template; scale bars represent 10 mm. A common t-statistic scale is used across Figures 2–4.

**Figure 3:**
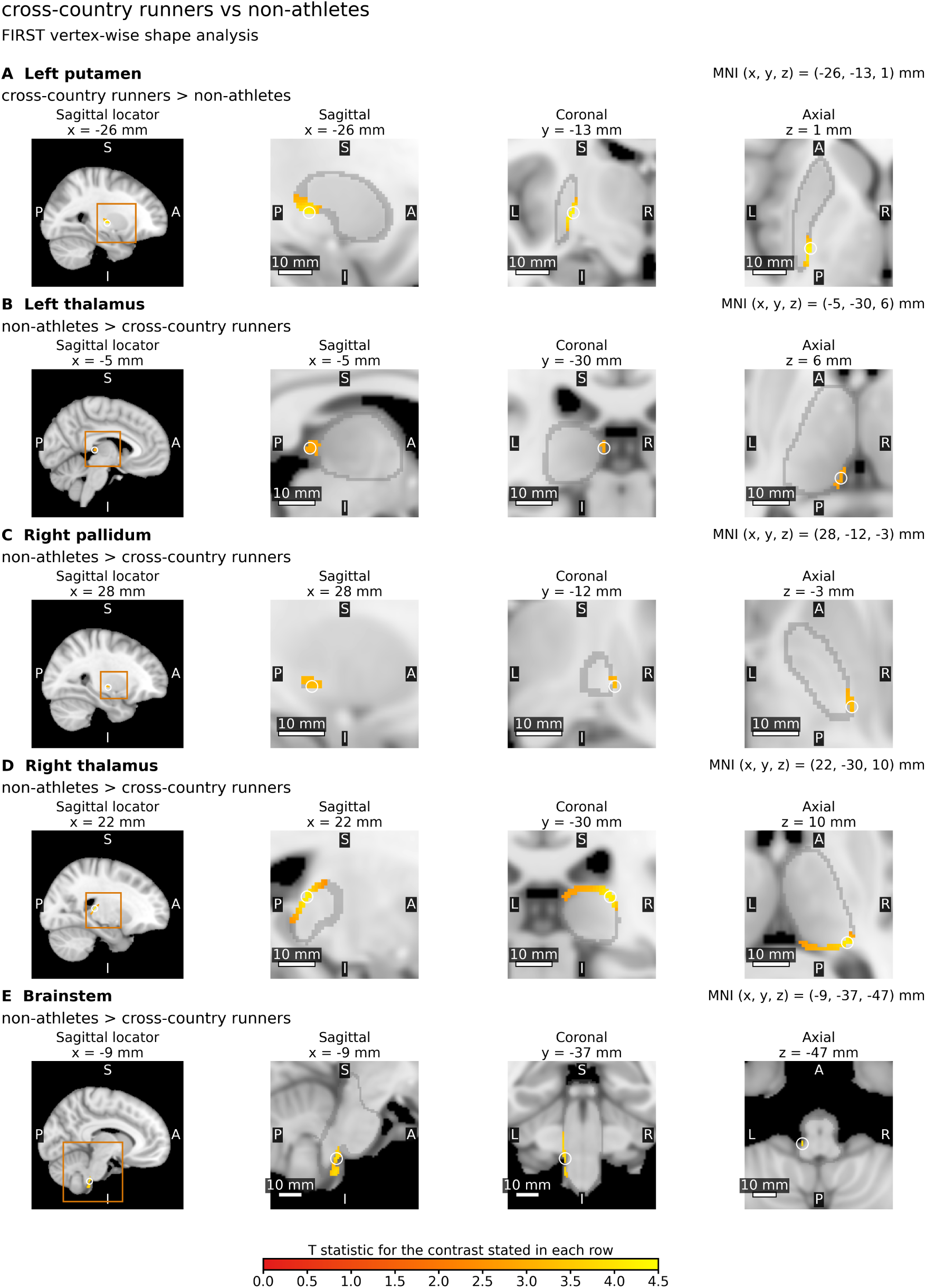
FIRST vertex-wise shape differences between cross-country runners and non-athletes. Significant local surface differences were identified in the left putamen (A), left thalamus (B), right pallidum (C), right thalamus (D), and brainstem (E). The left putamen shows the cross-country runners > non-athletes contrast, whereas the remaining structures show the non-athletes > cross-country runners contrast. Red-to-yellow overlays show directional t statistics at vertices surviving TFCE-based family-wise error correction (pFWE *≤* .05; 10,000 permutations). Each row shows a sagittal locator together with enlarged sagittal, coronal, and axial views intersecting the reported Montreal Neurological Institute (MNI) coordinate. Orange boxes in the locator images indicate the field of view of the enlarged sagittal view. Gray outlines delineate the tested FIRST structure, and white circles mark the reported MNI coordinate. Images are displayed on the MNI152 1-mm T1-weighted template; scale bars represent 10 mm. A common t-statistic scale is used across Figures 2–4.

**Figure 4:**
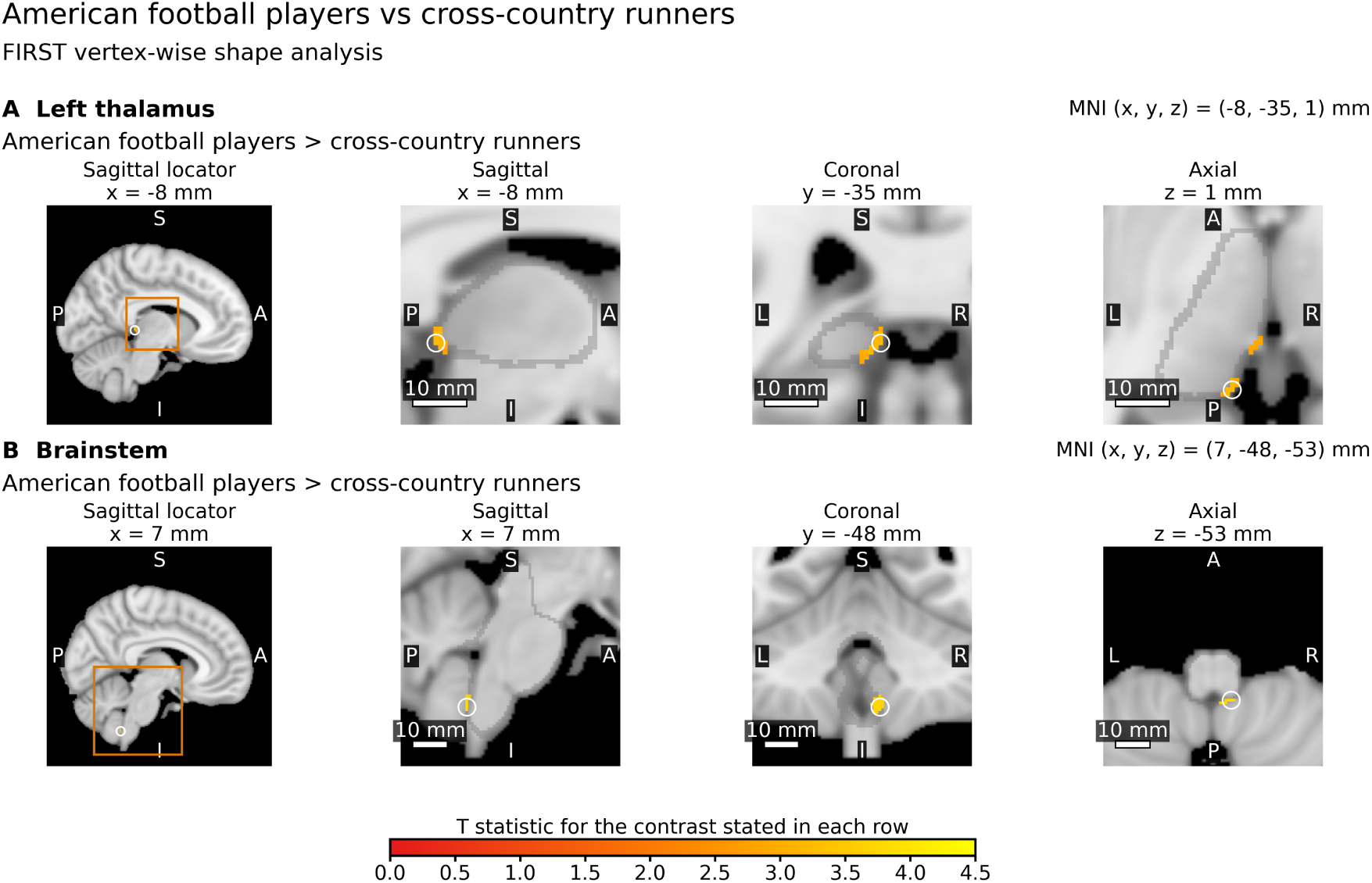
FIRST vertex-wise shape differences between American football players and cross-country runners. Significant local surface differences were identified in the left thalamus (A) and brainstem (B), both for the American football players > cross-country runners contrast. Red-to-yellow overlays show directional t statistics at vertices surviving TFCE-based family-wise error correction (pFWE *≤* .05; 10,000 permutations). Each row shows a sagittal locator together with enlarged sagittal, coronal, and axial views intersecting the reported Montreal Neurological Institute (MNI) coordinate. Orange boxes in the locator images indicate the field of view of the enlarged sagittal view. Gray outlines delineate the tested FIRST structure, and white circles mark the reported MNI coordinate. Images are displayed on the MNI152 1-mm T1-weighted template; scale bars represent 10 mm. A common t-statistic scale is used across Figures 2–4.

**Table 1:** Significant FIRST vertex-wise shape clusters from pairwise group comparisons. Vertex-wise shape analyses were performed on FIRST boundary-displacement maps using nonparametric permutation testing with TFCE correction. The table lists clusters surviving peak family-wise error correction at pFWE *≤* 0.05 in directional T-contrasts. Positive or negative effect-size signs follow the group-order coding used in the corresponding analysis rather than the verbal direction of every contrast. CI = confidence interval; MNI = Montreal Neurological Institute coordinate space; FWE = family-wise error; vox = voxels. Atlas locations are from the Harvard-Oxford Subcortical Structural Atlas.

| Comparison | Direction | Structure | Peak T | p-FWE | Cluster size (vox) | Cohen d | Hedges g | 95% CI | MNI x,y,z | Atlas location |
| --- | --- | --- | --- | --- | --- | --- | --- | --- | --- | --- |
| American football players vs non-athletes | non-athletes > American football players | Left nucleus accumbens | 4.07 | 0.0160 | 45 | -1.42 | -1.37 | [-2.32, -0.52] | -5,10,-4 | 42.1% Left Accumbens; 26.2% Left Cerebral White Matter; 12.0% Left Cerebral Cortex |
| American football players vs non-athletes | non-athletes > American football players | Right hippocampus | 4.30 | 0.0087 | 224 | -1.77 | -1.71 | [-2.72, -0.82] | 31,-9,-26 | 48.1% Right Hippocampus; 27.8% Right Cerebral White Matter; 4.5% Right Cerebral Cortex |
| American football players vs non-athletes | non-athletes > American football players | Brainstem | 4.20 | 0.0020 | 680 | -1.68 | -1.62 | [-2.61, -0.74] | -9,-37,-43 | 61.5% Brainstem |
| cross-country runners vs non-athletes | cross-country runners > non-athletes | Left putamen | 4.28 | 0.0127 | 186 | 1.59 | 1.54 | [0.66, 2.53] | -26,-13,1 | 50.2% Left Pallidum; 49.0% Left Putamen; 0.8% Left Cerebral White Matter |
| cross-country runners vs non-athletes | non-athletes > cross-country runners | Left thalamus | 2.92 | 0.0461 | 30 | -1.20 | -1.16 | [-2.09, -0.31] | -5,-30,6 | 62.0% Left Thalamus; 7.0% Left Cerebral White Matter |
| cross-country runners vs non-athletes | non-athletes > cross-country runners | Right pallidum | 3.31 | 0.0429 | 23 | -1.34 | -1.30 | [-2.25, -0.44] | 28,-12,-3 | 55.3% Right Putamen; 36.4% Right Pallidum; 8.3% Right Cerebral White Matter |
| cross-country runners vs non-athletes | non-athletes > cross-country runners | Right thalamus | 4.19 | 0.0079 | 416 | -1.42 | -1.37 | [-2.33, -0.51] | 22,-30,10 | 76.0% Right Cerebral White Matter; 23.8% Right Thalamus; 0.2% Right Lateral Ventricle |
| cross-country runners vs non-athletes | non-athletes > cross-country runners | Brainstem | 4.05 | 0.0323 | 84 | -1.88 | -1.81 | [-2.86, -0.90] | -9,-37,-47 | 55.7% Brainstem |
| American football players vs cross-country runners | American football players > cross-country runners | Left thalamus | 3.25 | 0.0355 | 120 | 1.16 | 1.13 | [0.41, 1.90] | -8,-35,1 | 43.8% Left Thalamus; 4.9% Left Cerebral White Matter; 1.1% Left Hippocampus |
| American football players vs cross-country runners | American football players > cross-country runners | Brainstem | 3.81 | 0.0441 | 37 | 1.33 | 1.30 | [0.57, 2.09] | 7,-48,-53 | 47.1% Brainstem |

For American football players versus non-athletes, three corrected clusters were identified, all in the non-athletes > American football players direction. The left nucleus accumbens contained a 45-voxel cluster with peak *T* = 4.07, *p*FWE = .0160, Hedges’ *g* = *−*1.37, 95% CI [*−*2.32, *−*0.52], and peak MNI coordinates [*−*5, 10, *−*4]. A larger cluster was present in the right hippocampus, comprising 224 voxels, peak *T* = 4.30, *p*FWE = .0087, Hedges’ *g* = *−*1.71, 95% CI [*−*2.72, *−*0.82], at [31, *−*9, *−*26]. The largest corrected cluster in this comparison was located in the brainstem and contained 680 voxels, peak *T* = 4.20, *p*FWE = .0020, Hedges’ *g* = *−*1.62, 95% CI [*−*2.61, *−*0.74], at [*−*9, *−*37, *−*43]. These spatial effects are illustrated in Figure 2.

The cross-country runner versus non-athlete comparison produced five corrected vertex-wise clusters and included effects in both directions. Cross-country runners showed greater local displacement than non-athletes in a 186-voxel left putamen/pallidum cluster, peak *T* = 4.28, *p*FWE = .0127, Hedges’ *g* = 1.54, 95% CI [0.66, 2.53], at [*−*26, *−*13, 1]. Four clusters showed the opposite direction. A left thalamic cluster contained 30 voxels, peak *T* = 2.92, *p*FWE = .0461, Hedges’ *g* = *−*1.16, 95% CI [*−*2.09, *−*0.31], at [*−*5, *−*30, 6]. A right pallidum/putamen cluster comprised 23 voxels, peak *T* = 3.31, *p*FWE = .0429, Hedges’ *g* = *−*1.30, 95% CI [*−*2.25, *−*0.44], at [28, *−*12, *−*3]. The largest subcortical cluster in this comparison was in the right thalamic region, with 416 voxels, peak *T* = 4.19, *p*FWE = .0079, Hedges’ *g* = *−*1.37, 95% CI [*−*2.33, *−*0.51], at [22, *−*30, 10]. A further brainstem cluster contained 84 voxels, peak *T* = 4.05, *p*FWE = .0323, Hedges’ *g* = *−*1.81, 95% CI [*−*2.86, *−*0.90], at [*−*9, *−*37, *−*47]. These findings are illustrated in Figure 3.

The direct comparison between American football players and cross-country runners identified two corrected clusters, both in the American football players > cross-country runners direction. A left thalamic cluster contained 120 voxels, peak *T* = 3.25, *p*FWE = .0355, Hedges’ *g* = 1.13, 95% CI [0.41, 1.90], at [*−*8, *−*35, 1]. A second cluster was located in the brainstem and contained 37 voxels, peak *T* = 3.81, *p*FWE = .0441, Hedges’ *g* = 1.30, 95% CI [0.57, 2.09], at [7, *−*48, *−*53], as illustrated in Figure 4.

The corrected vertex-wise results therefore differed according to the comparison being made. The football-versus-non-athlete effects were confined to the non-athletes > football direction, whereas the runner-versus-non-athlete analysis contained local effects in both directions. The direct athlete comparison identified left thalamic and brainstem effects in the football > runner direction. The complete corrected cluster statistics are reported in Table 1. Descriptive cluster-level effect-size estimates are additionally shown in Supplementary Figure S2.

### 3.3 FIRST total subcortical volumes

The total-volume analysis produced a different statistical pattern from the vertex-wise shape analysis. Specifically, none of the 15 FIRST-derived structures survived Benjamini-Hochberg FDR correction in any of the three pairwise comparisons. Complete group means, Welch tests, uncorrected *p* values, FDR-adjusted *q* values, and Hedges’ *g* estimates are therefore reported in Supplementary Table S2.

The numerically strongest uncorrected differences occurred principally in the direct football-versus-runner comparison. Left putamen volume was lower in American football players (5021.2 *±* 434.7 mm^3^) than in cross-country runners (5546.0 *±* 521.3 mm^3^), Welch *t* = *−*3.13, *p* = .0039, FDR *q* = .0590, Hedges’ *g* = *−*1.07. Right putamen volume showed a similar uncorrected pattern, with means of 5133.4 *±* 383.9 mm^3^ and 5561.4 *±* 512.4 mm^3^, respectively, Welch *t* = *−*2.70, *p* = .0116, FDR *q* = .0870, Hedges’ *g* = *−*0.93. Right hippocampal volume was also numerically lower in football players than runners, 4577.4 *±* 521.9 mm^3^ versus 5072.8 *±* 692.7 mm^3^, Welch *t* = *−*2.31, *p* = .0286, FDR *q* = .1428, Hedges’ *g* = *−*0.79. None of these comparisons met the corrected *q* < .05 criterion. Thus, corrected local surface-shape differences were detectable with FIRST even though no corresponding total-structure volumetric comparison survived FDR correction.

### 3.4 FAST whole-brain tissue volumes

FAST-derived tissue-volume distributions are illustrated in Figure 5. Table 2 summarizes the omnibus results from the primary unadjusted analysis and the eTIV-adjusted sensitivity analysis, while complete group-wise descriptive statistics are provided in Supplementary Table S3.

**Figure 5:**
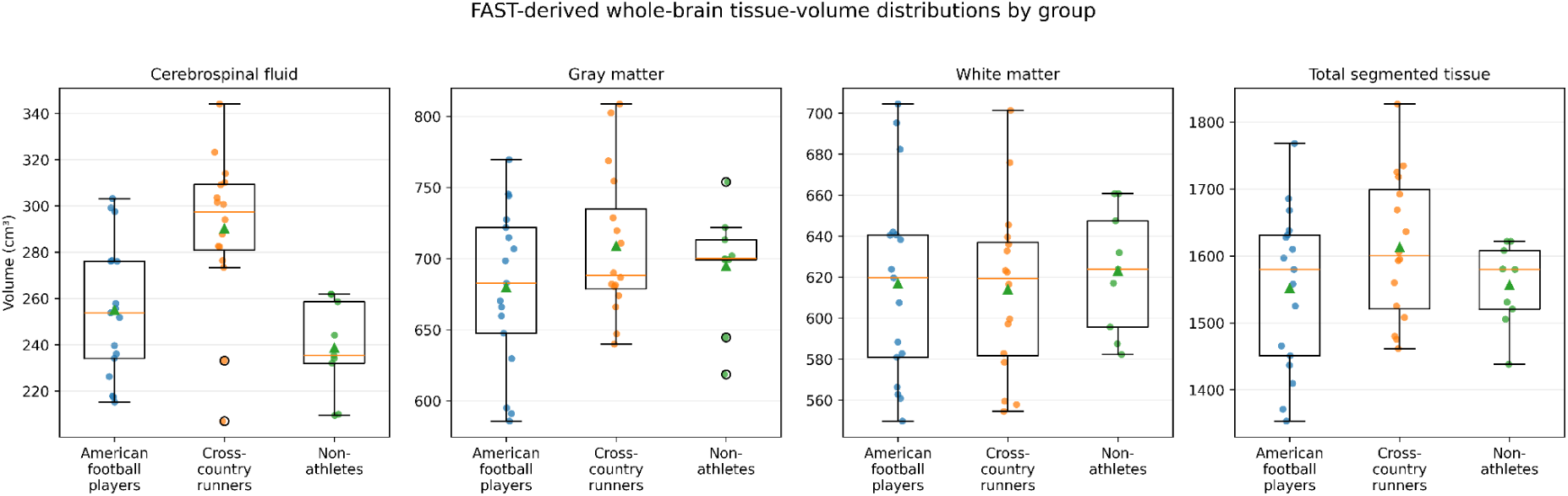
FAST-derived whole-brain tissue volumes by group. Box-and-whisker plots show cerebrospinal fluid (CSF), gray matter (GM), white matter (WM), and total segmented tissue volumes, expressed in cm^3^, for American football players (n=17), cross-country runners (n=16), and non-athletes (n=9). Total segmented tissue volume was calculated as CSF + GM + WM. Boxes indicate the interquartile range, center lines denote the median, whiskers extend to 1.5×IQR, the mean is indicated by the plotted mean marker, and jittered points represent individual participants.

**Table 2:** Omnibus group comparisons of FAST-derived tissue volumes before and after eTIV adjustment.

| Tissue-volume measure | Analysis | Omnibus statistic | *p* value | FDR *q* value |
| --- | --- | --- | --- | --- |
| **Panel A. Primary unadjusted analysis** |  |  |  |  |
| Cerebrospinal fluid volume | Welch ANOVA | $F(2.00, 24.38) = 11.46$ | $< .001$ | .001 |
| Gray matter volume | Welch ANOVA | $F(2.00, 23.51) = 1.16$ | .330 | .440 |
| White matter volume | Welch ANOVA | $F(2.00, 24.31) = 0.20$ | .820 | .820 |
| Total segmented tissue volume | Welch ANOVA | $F(2.00, 25.50) = 1.63$ | .215 | .430 |
| **Panel B. eTIV-adjusted sensitivity analysis** |  |  |  |  |
| Cerebrospinal fluid volume | HC3-robust Wald test | $F(2, 38) = 14.30$ | $< .001$ | $< .001$ |
| Gray matter volume | HC3-robust Wald test | $F(2, 38) = 0.08$ | .923 | .923 |
| White matter volume | HC3-robust Wald test | $F(2, 38) = 9.59$ | $< .001$ | $< .001$ |
| Total segmented tissue volume | HC3-robust Wald test | $F(2, 38) = 0.51$ | .604 | .806 |

*Note.* Panel A reports the primary unadjusted Welch analysis of variance across American football players, cross-country runners, and non-athletes. Panel B reports the head-size sensitivity analysis in which each FAST-derived absolute tissue volume was modeled as a function of group and mean-centered FreeSurfer estimated total intracranial volume (eTIV); the omnibus group effect was evaluated using a joint HC3 heteroskedasticity-robust Wald $F$ test. eTIV was centered at the sample mean of 1703.345 cm<sup>3</sup>. Benjamini-Hochberg false discovery rate (FDR) correction was applied separately across the four omnibus tests in Panel A and the four omnibus tests in Panel B. $p$ values and FDR $q$ values are two-sided.

In the unadjusted analysis, mean CSF volume was 254.9 *±* 29.2 cm^3^ in American football players, 290.2 *±* 33.2 cm^3^ in cross-country runners, and 238.6 *±* 20.2 cm^3^ in non-athletes. The three-group Welch ANOVA identified a group effect for CSF, *F* (2.00, 24.38) = 11.46, *p* < .001, FDR *q* = .001. By contrast, the corresponding omnibus tests did not meet the corrected criterion for gray matter, *F* (2.00, 23.51) = 1.16, *p* = .330, FDR *q* = .440; white matter, *F* (2.00, 24.31) = 0.20, *p* = .820, FDR *q* = .820; or total segmented tissue volume, *F* (2.00, 25.50) = 1.63, *p* = .215, FDR *q* = .430.

The corrected unadjusted pairwise CSF comparisons localized the omnibus effect primarily to the cross-country runner group. Cross-country runners had greater CSF volume than non-athletes by 51.59 cm^3^, 95% CI [29.49, 73.70], Welch *t* (22.76) = 4.83, *p* < .001, FDR *q* < .001, Hedges’ *g* = 1.70. American football players had lower CSF volume than cross-country runners, with a football-minus-runner mean difference of *−*35.26 cm^3^, 95% CI [*−*57.55, *−*12.97], Welch *t* (29.94) = *−*3.23, *p* = .003, FDR *q* = .018, Hedges’ *g* = *−*1.10. The football-versus-non-athlete CSF comparison did not survive FDR correction, mean difference = 16.33 cm^3^, 95% CI [*−*3.93, 36.60], Welch *t* (22.05) = 1.67, *p* = .109, FDR *q* = .264, Hedges’ *g* = 0.60.

None of the unadjusted pairwise comparisons for gray matter, white matter, or total segmented tissue volume survived FDR correction. Mean white matter volume was 616.8 *±* 48.0 cm^3^ in football players, 613.9 *±* 42.0 cm^3^ in runners, and 623.0 *±* 30.1 cm^3^ in non-athletes. Total segmented tissue volume was 1551.5 *±* 119.3, 1612.9 *±* 108.3, and 1556.3 *±* 62.1 cm^3^, respectively. Complete unadjusted pairwise results are provided in Supplementary Table S4.

Adjustment for eTIV altered part of the FAST pattern while retaining the corrected CSF group effect. Homogeneity of regression slopes was examined before the adjusted analysis, and no group-by-eTIV interaction was identified for any of the four tissue outcomes (all *p ≥* .349). Common-slope models were therefore used. In the eTIV-adjusted models with HC3 heteroskedasticity-robust inference, a corrected group effect remained for CSF, *F* (2, 38) = 14.30, *p* < .001, FDR *q* < .001. A corrected adjusted group effect was also present for white matter, *F* (2, 38) = 9.59, *p* < .001, FDR *q* < .001. By contrast, gray matter, *F* (2, 38) = 0.08, *p* = .923, FDR *q* = .923, and total segmented tissue volume, *F* (2, 38) = 0.51, *p* = .604, FDR *q* = .806, did not show corrected adjusted group effects.

At the grand mean eTIV of 1703.345 cm^3^, cross-country runners had greater adjusted CSF volume than non-athletes, adjusted mean difference = 41.52 cm^3^, 95% CI [24.70, 58.34], *t* (38) = 5.00, *p* < .001, FDR *q* < .001. American football players also had greater adjusted CSF volume than non-athletes, adjusted mean difference = 20.73 cm^3^, 95% CI [6.64, 34.82], *t* (38) = 2.98, *p* = .005, FDR *q* = .015. The adjusted American football player-versus-runner contrast was nominally different, adjusted mean difference = *−*20.79 cm^3^, 95% CI [*−*40.16, *−*1.43], *t* (38) = *−*2.17, *p* = .036, but did not survive FDR correction, *q* = .086.

For white matter, cross-country runners had lower adjusted volume than non-athletes, adjusted mean difference = *−*27.29 cm^3^, 95% CI [*−*40.58, *−*14.00], *t* (38) = *−*4.16, *p* < .001, FDR *q* = .001. American football players also had greater adjusted white matter volume than cross-country runners, adjusted mean difference = 29.05 cm^3^, 95% CI [11.29, 46.80], *t* (38) = 3.31, *p* = .002, FDR *q* = .008. The adjusted football-versus-non-athlete white matter contrast did not survive correction, *t* (38) = 0.22, *p* = .824, FDR *q* = .880. No corrected adjusted pairwise differences were identified for gray matter or total segmented tissue volume. Complete eTIV-adjusted pairwise results are provided in Supplementary Table S5. Descriptive tissue fractions are shown separately in Supplementary Figure S3.

### 3.5 Convergence across structural measures

Considered together, the three analyses showed both convergence and dissociation across measurement scales. First, the brainstem was repeatedly identified in the two spatial analyses. VBM showed substantial corrected brainstem clusters in both athlete-versus-non-athlete sets of balanced comparisons and in the direct football-versus-runner comparison, while FIRST vertex analysis independently identified corrected brainstem clusters for football versus non-athletes (*T* = 4.20, *p*FWE = .0020), runners versus non-athletes (*T* = 4.05, *p*FWE = .0323), and football versus runners (*T* = 3.81, *p*FWE = .0441). Second, the FIRST results differed according to measurement type: several localized vertex-wise effects survived FWE correction, whereas none of the 15 total-structure volume comparisons survived FDR correction. Third, the global FAST analysis showed a pattern that was partly sensitive to adjustment for intracranial size. The unadjusted analysis identified a corrected omnibus effect restricted to CSF, whereas the eTIV-adjusted sensitivity analysis retained the CSF group effect, *F* (2, 38) = 14.30, FDR *q* < .001, and additionally identified a white matter group effect, *F* (2, 38) = 9.59, FDR *q* < .001. Gray matter and total segmented tissue volume remained without corrected adjusted group effects. Thus, the global tissue findings differed according to whether absolute volumes or volumes adjusted for estimated intracranial size were considered.

## 4 Discussion

### 4.1 Principal findings

The study here examined structural brain differences among collegiate American football players, cross-country runners, and non-athletes using complementary measures of local gray matter morphology, subcortical volume and shape, and global tissue composition. Four findings are of primary relevance. First, VBM and FIRST provided converging evidence for brainstem-centered group differences, although the direction of these effects depended on the comparison being made. Non-athletes showed greater local measures than both athlete groups in the athlete-versus-non-athlete comparisons, whereas American football players showed greater brainstem measures than cross-country runners in the direct athlete comparison. Second, FIRST identified localized shape differences involving the nucleus accumbens, hippocampus, thalamus, basal ganglia, and brainstem, while none of the 15 FIRST-derived total structure volumes survived FDR correction at *q* < .05. Third, the cross-country runners showed a structural profile that was itself distinguishable from the non-athlete group, indicating that this group should not be treated simply as a neutral control for football exposure. Finally, the unadjusted FAST analysis identified a corrected CSF group effect, whereas gray matter, white matter, and total segmented tissue volume did not show corrected unadjusted effects. Adjustment for FreeSurfer eTIV retained the CSF group effect and additionally identified a corrected white matter group effect, while gray matter and total segmented tissue volume remained without corrected effects. The global tissue findings were therefore partly sensitive to adjustment for intracranial size. Taken together, the findings indicate structural variation across groups, but not a simple pattern of generalized increase or decrease associated specifically with American football.

### 4.2 Brainstem convergence across VBM and FIRST

The most consistent anatomical differences across the spatial analyses were in the brainstem. In the balanced VBM comparisons, non-athletes showed greater modulated gray matter volume than American football players in brainstem clusters in both athlete subsets. The same direction of effects was present when non-athletes were compared with cross-country runners. The direct athlete comparison, however, showed the opposite relative ordering: American football players showed greater modulated gray matter volume than cross-country runners in a brainstem cluster. FIRST provided a second form of evidence involving the same broad anatomical region, with corrected brainstem shape effects observed across all pairwise comparisons.

This convergence is noteworthy because VBM and FIRST characterize morphology differently. VBM tests local variation in spatially normalized and modulated gray matter images, whereas the FIRST vertex analysis tests localized surface displacement within a model-based representation of the brainstem and subcortical structures. Convergence across these analytical representations therefore makes the recurring brainstem localization more informative than a finding arising from only one measurement approach. It should not, however, be treated as an independent replication, because both analyses were derived from the same underlying T1-weighted images and can share sensitivity to image quality, registration, and anatomy.

More importantly, the direction of the findings argues against describing the brainstem result as a straightforward football-specific injury pattern. If repetitive football exposure alone were sufficient to explain the finding, a simple expectation might be that football players would differ consistently in the same direction from both comparison groups. This was not observed. Both football players and runners differed from non-athletes, while football players also differed from runners. This pattern leaves several possibilities open, including general characteristics associated with collegiate athletic participation, sport-specific training demands, pre-existing anatomical differences, collision-related exposure, or combinations of these factors.

Prior longitudinal studies make exposure-related explanations biologically plausible without establishing such an explanation for the current results. For example, MRI changes have been reported across a football season in athletes without clinically diagnosed concussion, including associations between head-impact exposure and white matter measures (Bazarian et al., 2014; Slobounov et al., 2017). Those studies differ from the present cross-sectional morphometric analysis and do not demonstrate that the brainstem effects reported here represent injury. Taken together, these findings indicate that brainstem morphology differed across analytical scales, while the origin and biological significance of these differences remains unresolved.

### 4.3 Subcortical shape differences without corrected volume differences

The FIRST results provide a second important distinction between local morphology and total structure size. In the football-versus-non-athlete comparison, corrected vertex-wise differences were identified in the left nucleus accumbens, the right hippocampus, and the brainstem. The runner-versus-non-athlete comparison produced corrected effects involving the putamen/pallidum, thalamus and brainstem, with effects occurring in both directions. The direct athlete comparison additionally identified a left thalamic effect in the football > runner direction. In contrast, none of the 15 total FIRST volumes survived FDR correction in any pairwise comparison. Even the largest uncorrected football-versus-runner volumetric difference did not meet the corrected criterion.

This distinction matters because surface shape and total volume are related but are not interchangeable anatomical measures. FIRST represents structures using model-based surface geometry, allowing localized boundary displacement to be identified even when the integrated volume of the complete structure is relatively similar between groups. A structure can therefore differ locally at particular parts of its surface without there being a difference in overall volume. For example, local outward displacement in one portion of a structure may also coexist with little difference, or even displacement in another direction, elsewhere on the structure.

At a systems level, the structures implicated here span cortico-subcortical networks rather than pointing toward a difference in one isolated anatomical system. The thalamus and basal ganglia contribute to distributed sensorimotor and associative circuits, while hippocampal and accumbens regions participate in learning-, memory-, and motivationally relevant networks. However, the presence of a local morphological difference does not establish altered function within those systems. No cognitive, clinical, or motor outcomes were available that could link the anatomical findings to performance or impairment. The finding should therefore be restricted to what was measured: localized surface morphology differed between groups even though corrected total structure volume did not.

This shape-volume dissociation also argues against reducing the FIRST findings to generalized enlargement or shrinkage of subcortical structures. At the same time, vertex-wise analyses remain sensitive to segmentation, registration, local image contrast, and individual anatomical variability. The shape findings are consequently most useful as evidence that the group differences were spatially localized within structures, rather than as evidence for a particular biological process.

### 4.4 The importance of the cross-country runner comparison

The inclusion of cross-country runners changes the interpretation substantially. In a football-versus-non-athlete design alone, differences between the groups could readily be framed around collision exposure. The present three-group design shows why that interpretation would be incomplete. Cross-country runners also differed from non-athletes in the VBM analysis, with both balanced contrasts showing brainstem effects. FIRST similarly identified runner-versus-non-athlete differences in multiple subcortical regions, including the brainstem, with effects not uniformly directed toward either larger or smaller local measures in runners.

The FAST analyses further reinforce the importance of treating the runner group as an active comparison population. In the unadjusted analysis, cross-country runners showed greater CSF volume than non-athletes and American football players. Adjustment for eTIV modified this pattern. The runner-versus-non-athlete CSF difference remained after correction, whereas the adjusted football-versus-runner CSF contrast did not survive FDR correction. American football players also showed greater adjusted CSF volume than non-athletes. In addition, the eTIV-adjusted analysis identified lower white matter volume in runners than non-athletes and American football players. Thus, the global tissue profile of the endurance-athlete group remained distinguishable from the other groups, but the particular pattern depended in part on whether differences in intracranial size were taken into account.

There is independent evidence that long-term endurance training can be associated with structural and functional brain differences. Cao et al. (2021), for example, reported structural and functional MRI differences between endurance runners and non-running controls. While the particular findings of that study cannot be used to explain the regions observed here directly, they do support the broader point that endurance athletes cannot be assumed to provide an anatomically neutral comparison population.

The runner group should therefore be viewed as an active comparison group with its own training history and physiological characteristics. This is one of the main advantages of the present design. Differences shared by both athlete groups weaken a uniquely collision-specific explanation, likely indicative of sport or fitness-related effects, whereas football-versus-runner differences identify patterns that may be more sport-specific. Even those differences, however, remain associations because the groups were not randomized for their sporting histories. In other words, differences between individuals likely relate to whether they choose to pursue any sport as well as the sport that they select. This is a notably under considered factor in many studies looking at effects of sport on cognition or brain structure.

### 4.5 Interpretation of the FAST findings

The FAST findings require particularly cautious interpretation because the statistical pattern differed between the unadjusted and eTIV-adjusted analyses. In the unadjusted analysis, CSF was the only tissue-volume measure showing a corrected omnibus group effect. Of relevance here, the runner-versus-non-athlete CSF difference remained after adjustment for FreeSurfer eTIV, indicating that this contrast was not accounted for solely by differences in estimated intracranial size. However, adjustment changed the organization of the pairwise CSF findings. American football players also showed greater adjusted CSF volume than non-athletes, whereas the football-versus-runner contrast no longer survived FDR correction. This means that the CSF finding should not be reduced to a simple runner-specific increase. More importantly, greater CSF volume should not itself be interpreted as evidence of, for example, cerebral atrophy, particularly because neither gray matter nor total segmented tissue volume showed a corresponding corrected adjusted reduction.

The eTIV-adjusted analysis also identified a white matter group effect that was not apparent in the unadjusted absolute-volume analysis. Cross-country runners showed lower adjusted white matter volume than non-athletes and American football players, whereas the football-versus-non-athlete comparison remained uncorrected. This difference is important because absolute white matter volume was strongly related to intracranial size in the present sample. Adjustment for eTIV therefore asks a different question from the unadjusted comparison: whether white matter volume differs between groups when participants are compared at a common estimated intracranial size. The finding should consequently be interpreted as a head-size-adjusted volumetric difference rather than evidence of white matter loss. Because the effect became apparent only after covariate adjustment and was identified within a sensitivity analysis, its biological meaning remains uncertain and requires independent replication.

The eTIV-adjusted findings therefore indicate that the FAST pattern is partly sensitive to anatomical scaling, while the biological meaning of these adjusted differences remains uncertain. In particular, the persistence of the runner-versus-non-athlete CSF difference after adjustment suggests that estimated intracranial size alone does not account for this contrast, whereas the altered CSF pairwise pattern and emergence of an adjusted white matter effect demonstrate that conclusions based on absolute tissue volumes can change when head-size variation is taken into account. These findings should therefore remain descriptive of group-level structural variation rather than being interpreted as evidence of a particular beneficial or adverse biological process.

### 4.6 Strengths and limitations

Several features strengthen the contribution of the study. The analysis extends a publicly available neuroimaging resource, allowing new structural questions to be addressed using data that can be independently accessed and reanalyzed. The original release was specifically intended to support further investigation of long-term collegiate athletic exposure and included 17 football players, 16 cross-country runners, and 9 non-athletes (Caron et al., 2021). More importantly, the design included both a non-athlete comparison group and a non-collision athlete group. This provides a more informative framework than a football-versus-non-athlete comparison alone because it allows differences associated with athletic participation more generally to be distinguished, at least descriptively, from differences that emerge between two athlete populations. A further strength is the use of complementary structural measures. VBM, FIRST shape, FIRST volumetry, and FAST do not answer the same anatomical question, and the brainstem convergence across VBM and FIRST is therefore informative even though these analyses share the same source images. In addition, the absolute FAST-derived tissue volumes were examined in a dedicated head-size sensitivity analysis using FreeSurfer eTIV values available from the public source dataset.

Several limitations restrict the conclusions that can be drawn. The sample was small for group-level structural neuroimaging, particularly the nine-person non-athlete group. Small samples increase uncertainty around effect estimates, sensitivity to individual observations, and the difficulty of incorporating potentially relevant covariates. This consideration also applies to the eTIV-adjusted FAST sensitivity analysis, which estimated group effects while additionally accounting for intracranial size. The adjusted findings, particularly the white matter effect that was not apparent in the unadjusted analysis, therefore require replication in a larger independent sample. The sample was also exclusively male and collegiate, meaning that the findings cannot be assumed to generalize to female athletes, younger players, former athletes, professional players, or athletes participating in other collision sports. The cross-sectional design is a further major limitation. It cannot establish whether the morphological differences preceded participation in a particular sport, emerged over years of training or exposure, or reflect some combination of selection and experience.

Exposure characterization was also limited. Football players with a diagnosed concussion within six months of the original study were excluded, and three retained football players had remote diagnosed concussions during college, but concussion history before college was unavailable. Detailed lifetime measures of head-impact frequency, magnitude, playing position, and cumulative exposure were not available. Consequently, the present analyses cannot test whether any structural measure varies in a dose-dependent manner with repetitive head-impact exposure. This limitation is particularly important because longitudinal football studies have shown that MRI measures can vary in association with impact exposure even in the absence of diagnosed concussion (Bazarian et al., 2014; Slobounov et al., 2017).

The secondary-analysis design also meant that structural findings could not be related to detailed cognitive, clinical, or behavioral outcomes. The source dataset did not contain the measures needed to determine whether individuals with particular morphological characteristics differed in memory, motor performance, symptoms, academic performance, or other functional outcomes. Structural group differences should therefore not be equated with functional benefit or impairment.

There are also analysis-specific limitations. The VBM analysis used balanced subset comparisons rather than one full 17/16/9 three-group model. This reduced sample-size imbalance for individual contrasts but meant that the VBM results represent a collection of balanced pairwise analyses rather than a single factorial test of all participants. The eTIV-adjusted FAST analysis also requires methodological qualification. FreeSurfer eTIV is an atlas-registration-based estimate rather than a direct segmentation of the intracranial cavity (Buckner et al., 2004), and previous methodological work has shown that eTIV can retain bias related to total brain volume (Klasson et al., 2018). The adjusted FAST findings should therefore be viewed as a head-size sensitivity analysis rather than definitive normalization of the tissue volumes. Finally, structural MRI measures remain sensitive to decisions concerning brain extraction, registration, smoothing, segmentation, and anatomical labeling. This issue is particularly relevant to small subcortical structures and the brainstem. Convergence across VBM and FIRST reduces dependence on a single analytical representation, but it does not eliminate sensitivity to preprocessing or anatomical variability.

### 4.7 Implications and future directions

The findings suggest that future work should move beyond simple collision-athlete-versus-control comparisons and combine better exposure characterization with repeated neuroimaging. Larger studies should include collision-sport athletes, non-collision athletes, and carefully characterized non-athletes, while also incorporating female participants and athletes across different ages and competitive levels. Longitudinal imaging before, during, and after periods of sporting exposure would be particularly valuable because it would allow within-participant structural trajectories to be separated from pre-existing group differences. Quantification of head impacts using exposure sensors or other validated measures, together with playing position, years of participation, concussion history, and training load, would make it possible to test whether structural outcomes vary with cumulative exposure rather than simply with group membership.

Future investigations should also connect anatomy with function. Cognitive, motor, symptom, and clinical measures would help determine whether local brainstem or subcortical morphology has behavioral relevance, rather than assuming that an anatomical difference necessarily represents benefit or harm. Multimodal integration is another priority. The public dataset already contains diffusion MRI derivatives, making it possible to examine whether gray matter morphology, subcortical shape, global tissue measures, and white matter microstructure show related participant-level patterns. Methodologically, replication of the eTIV-adjusted FAST pattern in a larger independent cohort would be particularly informative. Where possible, future studies could compare complementary methods of estimating intracranial size to determine whether the adjusted CSF and white matter patterns are stable across normalization approaches. Such work would help establish whether the head-size-sensitive findings reflect reproducible anatomical differences or are partly dependent on the particular intracranial-volume estimate used. Replication using a full-sample VBM model would also provide a useful extension of the present balanced-comparison approach. The current results therefore provide a basis for more targeted work, but the distinction between training-related adaptation, collision-related exposure, pre-existing anatomy, and normal individual variability will require longitudinal and multimodal evidence rather than structural MRI alone.

## 5 Conclusion

Structural MRI differences among American football players, cross-country runners, and non-athletes depend on both the comparison group and the anatomical scale being examined. Brainstem-centered effects were observed across VBM and FIRST shape analyses, but their direction was not uniquely associated with football, and localized FIRST shape differences occurred without corresponding FDR-corrected total subcortical-volume differences. Unadjusted FAST analyses identified a CSF group effect, while eTIV-adjusted sensitivity analyses showed that the CSF effect was retained and that white matter also differed across groups after accounting for estimated intracranial size; gray matter and total segmented tissue volume remained without corrected effects. The FAST pattern was therefore partly sensitive to anatomical scaling rather than being consistent with generalized brain-volume reduction. Overall, the findings do not support either a simple interpretation of generalized brain-volume loss associated with collegiate football or the assumption that all athlete-versus-non-athlete differences are football-specific. Longitudinal, exposure-informed, and independently replicated studies will be required to determine the origins and functional relevance of these structural patterns.

