## Supplementary Material for "Multiscale structural MRI differences among collegiate American football players, cross-country runners, and non-athletes"

---

#### S1. FIRST segmentation quality control

##### S1.1 Visual quality control of FIRST segmentations

All FIRST segmentation outputs were visually inspected before the volumetric and vertex-wise analyses. The purpose of this inspection was to verify that the model-derived subcortical labels corresponded plausibly to the underlying anatomy and that obvious segmentation or registration failures were not carried forward into the quantitative analyses. FIRST uses a model-based representation of subcortical shape and appearance, meaning that the anatomical plausibility of the resulting labels remains an important quality-control consideration before structure volumes or local surface displacement are evaluated (Patenaude et al., 2011).

Inspection focused on the correspondence between the segmentation boundaries and the visible anatomical margins of the bilateral nucleus accumbens, amygdala, caudate, hippocampus, pallidum, putamen, and thalamus, together with the brainstem. The segmentations were examined across sagittal, coronal, and axial views so that gross boundary displacement, misregistration, or extension of labels into anatomically implausible regions could be identified. No obvious segmentation failure requiring exclusion of an otherwise eligible participant was identified. Supplementary Figure S1 shows representative segmentation overlays from one participant in each group and is provided as an illustration of the quality-control procedure rather than as an outcome of the group analysis.

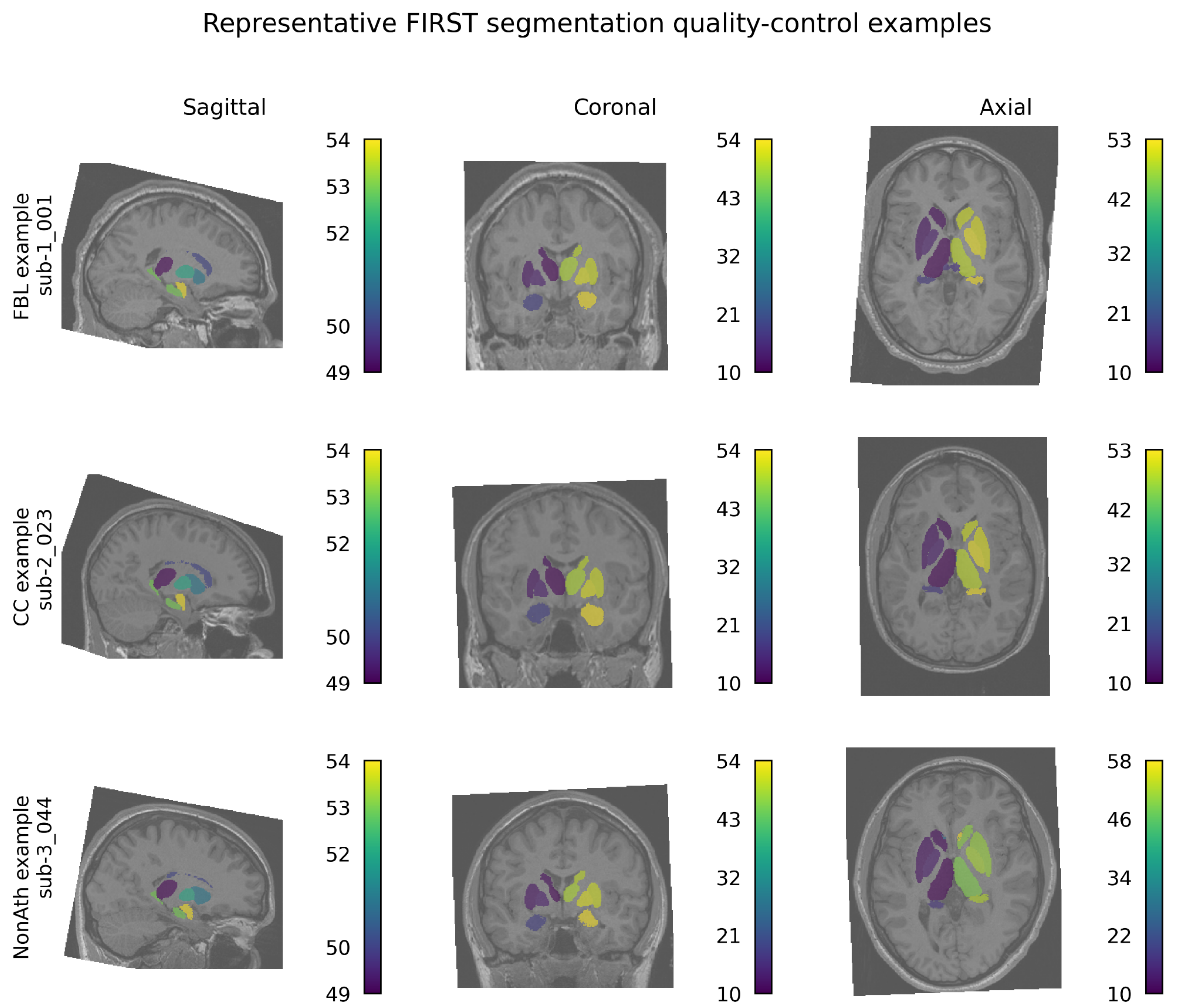


**Supplementary Figure S1:** FIRST segmentation quality-control examples. Representative FIRST segmentation overlays are shown for one American football player, one cross-country runner, and one non-athlete. For each participant, the FIRST-derived segmentation labels are overlaid on the corresponding T1-weighted anatomical image in sagittal, coronal, and axial views. The examples illustrate the anatomical correspondence and plausibility of the subcortical labels used for subsequent volume extraction and vertex-wise shape analysis. The displayed participants were selected as representative quality-control examples and were not selected on the basis of statistical outcome.

### S2. Complete voxel-based morphometry results

#### S2.1 Complete TFCE-corrected VBM cluster results

Supplementary Table S1 provides the complete cluster-level output from the balanced FSL-VBM comparisons reported in the main manuscript. The analyses comprised the first nine American football players versus the nine non-athletes, the last nine American football players versus the same nine non-athletes, the first nine cross-country runners versus the nine non-athletes, the last nine cross-country runners versus the nine non-athletes, and the balanced comparison of 16 American football players with 16 cross-country runners. Statistical inference was based on permutation testing with threshold-free cluster enhancement, which provides spatial inference without requiring selection of an initial cluster-forming threshold (Smith & Nichols, 2009).

**Supplementary Table S1:** Whole-brain significant clusters from FSL-VBM analyses. Significant clusters were obtained from TFCE-corrected permutation tests using FSL randomise with 5,000 permutations. Corrected $1 - p$ images were thresholded at 0.95, corresponding to corrected $p \leq .05$. The table reports the analysis and contrast, anatomical location, cluster extent (k), maximum corrected $1 - p$ value, corresponding TFCE-corrected p value, peak MNI coordinate, center-of-gravity MNI coordinate, maximum t statistic and its coordinate, and mean t statistic. Coordinates are reported in MNI152 2-mm space.

| Panel | Analysis | Contrast | Cluster | Anatomical location | k | 1-pmax | pTFCE-corr | Peak MNI x,y,z | COG MNI x,y,z | tmax | tmax MNI x,y,z | tmean |
|---|---|---|---|---|---|---|---|---|---|---|---|---|
| A | First nine American football players vs non-athletes | NonAth > FBL (tstat2) | 2 | Brain-Stem | 1,560 | 1.000 | <0.001 | 8, -38, -58 | 6.16, -39.7, -48.8 | 12.60 | 8, -38, -56 | 4.62 |
| A | First nine American football players vs non-athletes | NonAth > FBL (tstat2) | 1 | Right Cerebral Cortex | 3 | 0.958 | 0.042 | 28, 46, 24 | 26.7, 46.7, 24.0 | 7.37 | 28, 46, 24 | 7.27 |
| B | Last nine American football players vs non-athletes | NonAth > FBL (tstat2) | 2 | Brain-Stem | 1,165 | 0.995 | 0.005 | 8, -40, -58 | 0.92, -37.9, -48.8 | 6.73 | 8, -40, -58 | 4.45 |
| B | Last nine American football players vs non-athletes | NonAth > FBL (tstat2) | 1 | Right VIIIa | 9 | 0.955 | 0.045 | 38, -52, -60 | 37.3, -51.8, -58.4 | 6.32 | 38, -52, -60 | 5.76 |
| C | First nine cross-country runners vs non-athletes | NonAth > CC (tstat2) | 1 | Brain-Stem | 3,228 | 1.000 | <0.001 | 4, -40, -62 | 2.69, -42.1, -44.8 | 10.50 | 4, -36, -54 | 4.76 |
| D | Last nine cross-country runners vs non-athletes | NonAth > CC (tstat2) | 1 | Brain-Stem | 3,964 | 1.000 | <0.001 | 0, -42, -66 | 4.11, -39.9, -42.3 | 10.60 | 0, -42, -40 | 4.60 |
| E | Sixteen American football players vs sixteen cross-country runners | FBL > CC (tstat1) | 2 | Brain-Stem | 275 | 0.981 | 0.019 | 10, -42, -40 | 3.32, -38.1, -33.5 | 5.30 | 10, -42, -40 | 3.85 |
| E | Sixteen American football players vs sixteen cross-country runners | FBL > CC (tstat1) | 1 | Brain-Stem | 2 | 0.950 | 0.050 | -2, -44, -48 | -2.0, -44.0, -47.0 | 3.83 | -2, -44, -46 | 3.82 |

FBL = American football players; CC = cross-country runners; NonAth = non-athletes; COG = center of gravity; TFCE = threshold-free cluster enhancement; MNI = Montreal Neurological Institute.

### S3. Additional FIRST results

#### S3.1 Descriptive effect sizes for corrected vertex-wise clusters

Supplementary Figure S2 provides a standardized descriptive summary of the group differences within the FIRST vertex-wise clusters that survived TFCE-based family-wise error correction in the primary spatial analysis. Hedges' $g$ and the associated 95% confidence interval are displayed for each corrected cluster so that the direction and relative magnitude of the participant-level displacement differences can be considered alongside the inferential statistics reported in the main manuscript. These effect-size estimates are intended to characterize the corrected clusters rather than provide an additional statistical test.

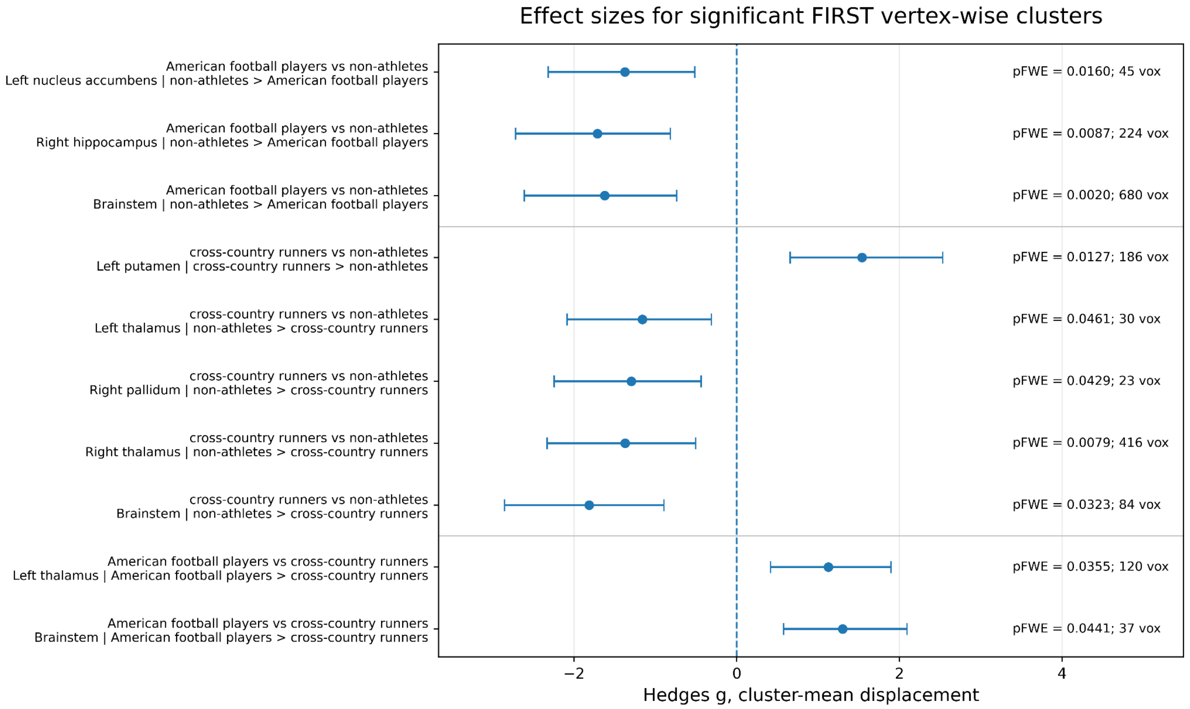


**Supplementary Figure S2:** Descriptive effect sizes for corrected FIRST vertex-wise clusters. Forest plot summarizing standardized effect sizes for the directional FIRST vertex-wise clusters surviving TFCE/FWE correction in the main analysis. Points represent cluster-mean Hedges' g, and horizontal bars represent 95% confidence intervals. Cluster-level pFWE values and cluster sizes are shown alongside the estimates. Effect-size signs follow the group-order coding used in each pairwise analysis: American football players minus non-athletes, cross-country runners minus non-athletes, and American football players minus cross-country runners. Accordingly, negative estimates in contrasts expressed as non-athletes > American football players or non-athletes > cross-country runners indicate greater cluster-mean displacement in the second-named group. FWE = family-wise error; vox = voxels.

##### S3.2 Complete FIRST total-volume analysis

Supplementary Table S2 provides the complete FIRST total-volume analysis for all 15 segmented structures. The table includes the bilateral nucleus accumbens, amygdala, caudate, hippocampus, pallidum, putamen, and thalamus, together with the brainstem, and reports the three pairwise comparisons used throughout the study: American football players versus non-athletes, cross-country runners versus non-athletes, and American football players versus cross-country runners.

For each structure, Supplementary Table S2 provides group descriptive statistics together with the corresponding Welch independent-samples test, uncorrected $p$ value, FDR-adjusted $q$ value, and standardized effect-size estimate. The Benjamini-Hochberg procedure was used to control the false discovery rate across the 15 structure-wise tests within each pairwise group comparison (Benjamini & Hochberg, 1995). Across the three comparison families, no FIRST total-volume comparison met the predefined FDR-corrected criterion of $q < .05$.

**Supplementary Table S2:** FIRST subcortical volume summaries and pairwise Welch tests. Descriptive volumes and pairwise Welch independent-samples tests are reported for the 15 FIRST-derived structures across American football players, cross-country runners, and non-athletes. Values include group mean ± SD, Welch t statistic, uncorrected p value, Benjamini-Hochberg FDR-adjusted q value, Cohen's d, and Hedges' g where available. FDR correction was applied across the 15 structures separately within each pairwise comparison. No comparison survived correction at q < .05. Volumes are expressed in $mm^3$. FDR = false discovery rate.

| Comparison | Structure | Group 1 | Group 1 mean ± SD (mm3) | Group 2 | Group 2 mean ± SD (mm3) | Welch t | p | FDR q | Hedges g | FDR significant |
|---|---|---|---|---|---|---|---|---|---|---|
| American football players vs non-athletes | Left nucleus accumbens | American football players (n=17) | 731.5 ± 193.4 | non-athletes (n=9) | 887.8 ± 128.4 | -2.46 | 0.0220 | 0.1717 | -0.87 | No |
| American football players vs non-athletes | Right nucleus accumbens | American football players (n=17) | 627.6 ± 162.7 | non-athletes (n=9) | 734.3 ± 79.4 | -2.24 | 0.0343 | 0.1717 | -0.73 | No |
| American football players vs non-athletes | Left amygdala | American football players (n=17) | 1757.9 ± 330.6 | non-athletes (n=9) | 1907.6 ± 336.2 | -1.09 | 0.2934 | 0.4881 | -0.44 | No |
| American football players vs non-athletes | Right amygdala | American football players (n=17) | 1663.3 ± 350.6 | non-athletes (n=9) | 1889.2 ± 229.3 | -1.98 | 0.0604 | 0.2267 | -0.69 | No |
| American football players vs non-athletes | Left caudate nucleus | American football players (n=17) | 4903.7 ± 518.9 | non-athletes (n=9) | 5234.9 ± 473.7 | -1.64 | 0.1185 | 0.2963 | -0.64 | No |
| American football players vs non-athletes | Right caudate nucleus | American football players (n=17) | 4957.4 ± 431.6 | non-athletes (n=9) | 5058.3 ± 640.5 | -0.42 | 0.6787 | 0.7831 | -0.19 | No |
| American football players vs non-athletes | Left hippocampus | American football players (n=17) | 4927.2 ± 614.9 | non-athletes (n=9) | 4680.2 ± 472.3 | 1.14 | 0.2679 | 0.4881 | 0.42 | No |
| American football players vs non-athletes | Right hippocampus | American football players (n=17) | 4577.4 ± 521.9 | non-athletes (n=9) | 4962.5 ± 343.8 | -2.26 | 0.0341 | 0.1717 | -0.79 | No |
| American football players vs non-athletes | Left pallidum | American football players (n=17) | 1814.1 ± 159.5 | non-athletes (n=9) | 1825.1 ± 75.1 | -0.24 | 0.8133 | 0.8504 | -0.08 | No |
| American football players vs non-athletes | Right pallidum | American football players (n=17) | 1802.6 ± 126.2 | non-athletes (n=9) | 1845.8 ± 89.4 | -1.01 | 0.3234 | 0.4881 | -0.36 | No |
| American football players vs non-athletes | Left putamen | American football players (n=17) | 5021.2 ± 434.7 | non-athletes (n=9) | 5289.1 ± 374.8 | -1.64 | 0.1180 | 0.2963 | -0.62 | No |
| American football players vs non-athletes | Right putamen | American football players (n=17) | 5133.4 ± 383.9 | non-athletes (n=9) | 5280.5 ± 335.8 | -1.01 | 0.3254 | 0.4881 | -0.39 | No |
| American football players vs non-athletes | Left thalamus | American football players (n=17) | 8938.2 ± 799.1 | non-athletes (n=9) | 8892.2 ± 428.6 | 0.19 | 0.8504 | 0.8504 | 0.06 | No |
| American football players vs non-athletes | Right thalamus | American football players (n=17) | 8652.9 ± 789.0 | non-athletes (n=9) | 8798.2 ± 319.8 | -0.66 | 0.5138 | 0.6703 | -0.21 | No |
| American football players vs non-athletes | Brainstem | American football players (n=17) | 25606.5 ± 3687.5 | non-athletes (n=9) | 26194.7 ± 803.7 | -0.63 | 0.5363 | 0.6703 | -0.19 | No |
| cross-country runners vs non-athletes | Left nucleus accumbens | cross-country runners (n=16) | 767.6 ± 146.6 | non-athletes (n=9) | 887.8 ± 128.4 | -2.13 | 0.0466 | 0.3753 | -0.83 | No |
| cross-country runners vs non-athletes | Right nucleus accumbens | cross-country runners (n=16) | 666.9 ± 127.0 | non-athletes (n=9) | 734.3 ± 79.4 | -1.63 | 0.1170 | 0.3753 | -0.58 | No |
| cross-country runners vs non-athletes | Left amygdala | cross-country runners (n=16) | 1714.0 ± 310.0 | non-athletes (n=9) | 1907.6 ± 336.2 | -1.42 | 0.1751 | 0.3753 | -0.59 | No |
| cross-country runners vs non-athletes | Right amygdala | cross-country runners (n=16) | 1768.7 ± 295.1 | non-athletes (n=9) | 1889.2 ± 229.3 | -1.13 | 0.2699 | 0.4049 | -0.43 | No |
| cross-country runners vs non-athletes | Left caudate nucleus | cross-country runners (n=16) | 4969.2 ± 680.8 | non-athletes (n=9) | 5234.9 ± 473.7 | -1.14 | 0.2648 | 0.4049 | -0.42 | No |
| cross-country runners vs non-athletes | Right caudate nucleus | cross-country runners (n=16) | 5146.1 ± 747.6 | non-athletes (n=9) | 5058.3 ± 640.5 | 0.31 | 0.7603 | 0.7882 | 0.12 | No |
| cross-country runners vs non-athletes | Left hippocampus | cross-country runners (n=16) | 5116.7 ± 561.9 | non-athletes (n=9) | 4680.2 ± 472.3 | 2.07 | 0.0522 | 0.3753 | 0.79 | No |
| cross-country runners vs non-athletes | Right hippocampus | cross-country runners (n=16) | 5072.8 ± 692.7 | non-athletes (n=9) | 4962.5 ± 343.8 | 0.53 | 0.6004 | 0.6928 | 0.18 | No |
| cross-country runners vs non-athletes | Left pallidum | cross-country runners (n=16) | 1886.6 ± 125.0 | non-athletes (n=9) | 1825.1 ± 75.1 | 1.54 | 0.1385 | 0.3753 | 0.54 | No |

*Continued on next page*

*Supplementary Table S2 (continued)*

| **Comparison** | **Structure** | **Group 1** | **Group 1 mean ± SD (mm3)** | **Group 2** | **Group 2 mean ± SD (mm3)** | **Welch t** | **p** | **FDR q** | **Hedges g** | **FDR significant** |
|---|---|---|---|---|---|---|---|---|---|---|
| cross-country runners vs non-athletes | Right pallidum | cross-country runners (n=16) | 1888.9 ± 176.8 | non-athletes (n=9) | 1845.8 ± 89.4 | 0.81 | 0.4268 | 0.5819 | 0.27 | No |
| cross-country runners vs non-athletes | Left putamen | cross-country runners (n=16) | 5546.0 ± 521.3 | non-athletes (n=9) | 5289.1 ± 374.8 | 1.42 | 0.1691 | 0.3753 | 0.52 | No |
| cross-country runners vs non-athletes | Right putamen | cross-country runners (n=16) | 5561.4 ± 512.4 | non-athletes (n=9) | 5280.5 ± 335.8 | 1.65 | 0.1127 | 0.3753 | 0.59 | No |
| cross-country runners vs non-athletes | Left thalamus | cross-country runners (n=16) | 9026.0 ± 645.5 | non-athletes (n=9) | 8892.2 ± 428.6 | 0.62 | 0.5411 | 0.6764 | 0.22 | No |
| cross-country runners vs non-athletes | Right thalamus | cross-country runners (n=16) | 8857.9 ± 768.1 | non-athletes (n=9) | 8798.2 ± 319.8 | 0.27 | 0.7882 | 0.7882 | 0.09 | No |
| cross-country runners vs non-athletes | Brainstem | cross-country runners (n=16) | 27028.1 ± 2725.3 | non-athletes (n=9) | 26194.7 ± 803.7 | 1.14 | 0.2690 | 0.4049 | 0.36 | No |
| American football players vs cross-country runners | Left nucleus accumbens | American football players (n=17) | 731.5 ± 193.4 | cross-country runners (n=16) | 767.6 ± 146.6 | -0.61 | 0.5482 | 0.6853 | -0.20 | No |
| American football players vs cross-country runners | Right nucleus accumbens | American football players (n=17) | 627.6 ± 162.7 | cross-country runners (n=16) | 666.9 ± 127.0 | -0.78 | 0.4440 | 0.6208 | -0.26 | No |
| American football players vs cross-country runners | Left amygdala | American football players (n=17) | 1757.9 ± 330.6 | cross-country runners (n=16) | 1714.0 ± 310.0 | 0.39 | 0.6966 | 0.7595 | 0.13 | No |
| American football players vs cross-country runners | Right amygdala | American football players (n=17) | 1663.3 ± 350.6 | cross-country runners (n=16) | 1768.7 ± 295.1 | -0.94 | 0.3562 | 0.6208 | -0.32 | No |
| American football players vs cross-country runners | Left caudate nucleus | American football players (n=17) | 4903.7 ± 518.9 | cross-country runners (n=16) | 4969.2 ± 680.8 | -0.31 | 0.7595 | 0.7595 | -0.11 | No |
| American football players vs cross-country runners | Right caudate nucleus | American football players (n=17) | 4957.4 ± 431.6 | cross-country runners (n=16) | 5146.1 ± 747.6 | -0.88 | 0.3871 | 0.6208 | -0.30 | No |
| American football players vs cross-country runners | Left hippocampus | American football players (n=17) | 4927.2 ± 614.9 | cross-country runners (n=16) | 5116.7 ± 561.9 | -0.93 | 0.3620 | 0.6208 | -0.31 | No |
| American football players vs cross-country runners | Right hippocampus | American football players (n=17) | 4577.4 ± 521.9 | cross-country runners (n=16) | 5072.8 ± 692.7 | -2.31 | 0.0286 | 0.1428 | -0.79 | No |
| American football players vs cross-country runners | Left pallidum | American football players (n=17) | 1814.1 ± 159.5 | cross-country runners (n=16) | 1886.6 ± 125.0 | -1.46 | 0.1554 | 0.4662 | -0.49 | No |
| American football players vs cross-country runners | Right pallidum | American football players (n=17) | 1802.6 ± 126.2 | cross-country runners (n=16) | 1888.9 ± 176.8 | -1.60 | 0.1202 | 0.4507 | -0.55 | No |
| American football players vs cross-country runners | Left putamen | American football players (n=17) | 5021.2 ± 434.7 | cross-country runners (n=16) | 5546.0 ± 521.3 | -3.13 | 0.0039 | 0.0590 | -1.07 | No |
| American football players vs cross-country runners | Right putamen | American football players (n=17) | 5133.4 ± 383.9 | cross-country runners (n=16) | 5561.4 ± 512.4 | -2.70 | 0.0116 | 0.0870 | -0.93 | No |
| American football players vs cross-country runners | Left thalamus | American football players (n=17) | 8938.2 ± 799.1 | cross-country runners (n=16) | 9026.0 ± 645.5 | -0.35 | 0.7300 | 0.7595 | -0.12 | No |
| American football players vs cross-country runners | Right thalamus | American football players (n=17) | 8652.9 ± 789.0 | cross-country runners (n=16) | 8857.9 ± 768.1 | -0.76 | 0.4552 | 0.6208 | -0.26 | No |
| American football players vs cross-country runners | Brainstem | American football players (n=17) | 25606.5 ± 3687.5 | cross-country runners (n=16) | 27028.1 ± 2725.3 | -1.26 | 0.2160 | 0.5400 | -0.43 | No |

### S4. Additional FAST results

#### S4.1 Descriptive whole-brain tissue volumes

Supplementary Table S3 provides the complete descriptive statistics underlying the FAST-derived tissue-volume distributions shown in main-text Figure 5. Values are reported separately for cerebrospinal fluid, gray matter, white matter, and total segmented tissue volume in American football players ($n = 17$), cross-country runners ($n = 16$), and non-athletes ($n = 9$). FAST uses probabilistic tissue classification and can produce partial-volume estimates for the principal tissue classes, allowing fractional tissue contributions at boundary voxels to contribute to volumetric estimation rather than requiring each voxel to be assigned exclusively to a single class (Zhang et al., 2001).

**Supplementary Table S3:** Whole-brain tissue volumes from FAST segmentation. Group-wise descriptive statistics are reported for FAST-derived cerebrospinal fluid, gray matter, white matter, and total segmented tissue volumes. Values are mean ± SD in $cm^3$. Tissue volumes were derived from the FAST partial-volume estimate maps and converted from $mm^3$ to $cm^3$ for presentation. Total segmented tissue volume was calculated as the sum of cerebrospinal fluid, gray matter, and white matter volumes. American football players, n = 17; cross-country runners, n = 16; non-athletes, n = 9.

| **Group** | **n** | **Cerebrospinal fluid volume ($cm^3$)** | **Gray matter volume ($cm^3$)** | **White matter volume ($cm^3$)** | **Total segmented tissue volume ($cm^3$)** |
|---|---|---|---|---|---|
| American football players | 17 | 254.9 ± 29.2 | 679.7 ± 56.5 | 616.8 ± 48.0 | 1551.5 ± 119.3 |
| Cross-country runners | 16 | 290.2 ± 33.2 | 708.8 ± 51.4 | 613.9 ± 42.0 | 1612.9 ± 108.3 |
| Non-athletes | 9 | 238.6 ± 20.2 | 694.7 ± 40.3 | 623.0 ± 30.1 | 1556.3 ± 62.1 |

#### S4.2 Complete unadjusted pairwise FAST comparisons

Supplementary Table S4 presents the complete pairwise inferential results for the four FAST-derived tissue-volume outcomes. The table includes American football players versus non-athletes, cross-country runners versus non-athletes, and American football players versus cross-country runners for cerebrospinal fluid, gray matter, white matter, and total segmented tissue volume. This provides the complete set of follow-up comparisons underlying the more selective presentation in the main Results, where emphasis is placed on the corrected cerebrospinal fluid comparisons that account for the omnibus group effect.

**Supplementary Table S4:** Group comparisons of FAST-derived tissue volumes: unadjusted pairwise group comparisons. Pairwise Welch independent-samples t tests are reported for cerebrospinal fluid, gray matter, white matter, and total segmented tissue volume. Mean differences are expressed as Group A minus Group B in $cm^3$ and are accompanied by 95% confidence intervals. Positive mean differences and positive Hedges' g values indicate greater volume in the first-named group. Benjamini-Hochberg FDR-adjusted q values were calculated across the full set of pairwise tissue-volume tests. CI = confidence interval; FDR = false discovery rate.

| Tissue-volume measure | Contrast | Mean difference ($cm^3$) [95% CI] | Welch t statistic | p value | FDR q value | Hedges g |
|---|---|---|---|---|---|---|
| Cerebrospinal fluid volume | American football players - Non-athletes | 16.33 [-3.93, 36.60] | t(22.05) = 1.67 | 0.109 | 0.264 | 0.60 |
| Cerebrospinal fluid volume | Cross-country runners - Non-athletes | 51.59 [29.49, 73.70] | t(22.76) = 4.83 | <0.001 | <0.001 | 1.70 |
| Cerebrospinal fluid volume | American football players - Cross-country runners | -35.26 [-57.55, -12.97] | t(29.94) = -3.23 | 0.003 | 0.018 | -1.10 |
| Gray matter volume | American football players - Non-athletes | -14.97 [-54.83, 24.88] | t(21.59) = -0.78 | 0.444 | 0.687 | -0.28 |
| Gray matter volume | Cross-country runners - Non-athletes | 14.08 [-24.70, 52.85] | t(20.28) = 0.76 | 0.458 | 0.687 | 0.28 |
| Gray matter volume | American football players - Cross-country runners | -29.05 [-67.37, 9.27] | t(30.97) = -1.55 | 0.132 | 0.264 | -0.52 |
| White matter volume | American football players - Non-athletes | -6.19 [-37.97, 25.58] | t(23.11) = -0.40 | 0.691 | 0.829 | -0.14 |
| White matter volume | Cross-country runners - Non-athletes | -9.09 [-39.26, 21.08] | t(21.44) = -0.63 | 0.538 | 0.717 | -0.23 |
| White matter volume | American football players - Cross-country runners | 2.90 [-29.10, 34.90] | t(30.85) = 0.18 | 0.855 | 0.893 | 0.06 |
| Total segmented tissue volume | American football players - Non-athletes | -4.83 [-78.27, 68.60] | t(24.00) = -0.14 | 0.893 | 0.893 | -0.05 |
| Total segmented tissue volume | Cross-country runners - Non-athletes | 56.58 [-13.96, 127.12] | t(22.96) = 1.66 | 0.111 | 0.264 | 0.58 |
| Total segmented tissue volume | American football players - Cross-country runners | -61.41 [-142.25, 19.43] | t(30.96) = -1.55 | 0.131 | 0.264 | -0.52 |

##### S4.3 Descriptive tissue-fraction composition

Supplementary Figure S3 provides a descriptive representation of tissue composition after each FAST-derived tissue volume was expressed relative to total segmented tissue volume.

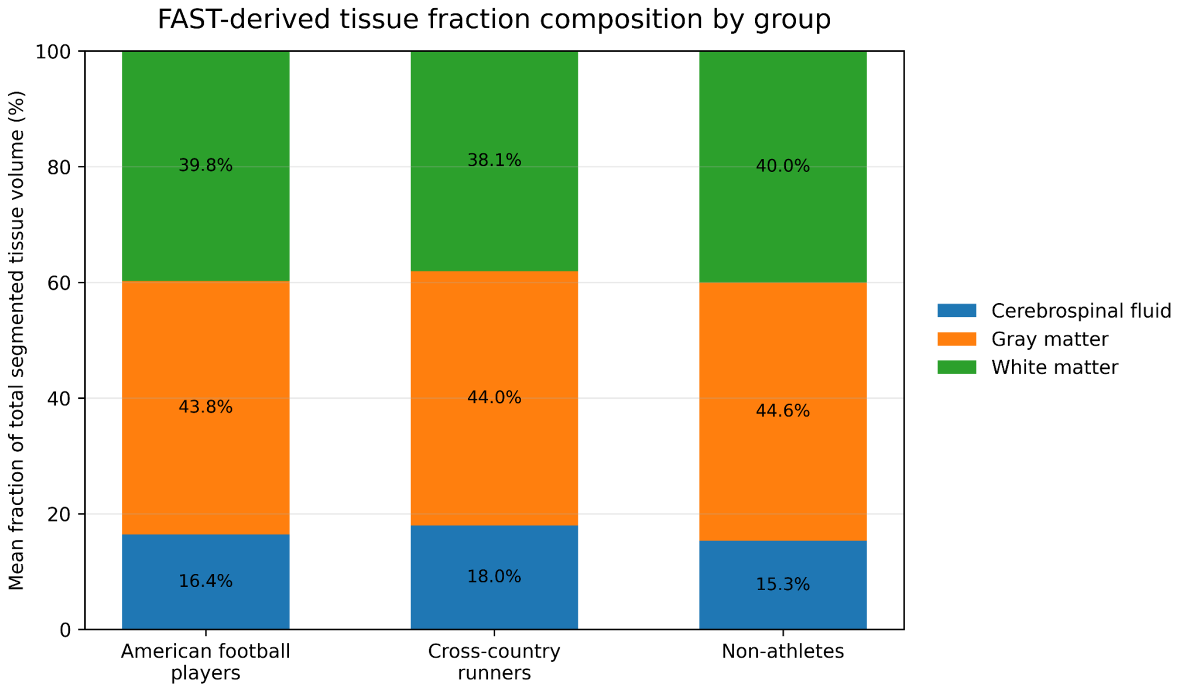


**Supplementary Figure S3:** Descriptive tissue-fraction composition. Stacked bars show the mean percentage contribution of cerebrospinal fluid, gray matter, and white matter to total segmented tissue volume in American football players, cross-country runners, and non-athletes. Tissue fractions were calculated as the volume of each tissue class divided by total segmented tissue volume and multiplied by 100. Group means were: American football players, cerebrospinal fluid 16.4%, gray matter 43.8%, white matter 39.8%; cross-country runners, cerebrospinal fluid 18.0%, gray matter 44.0%, white matter 38.1%; and non-athletes, cerebrospinal fluid 15.3%, gray matter 44.6%, white matter 40.0%.

#### S4.4 eTIV-adjusted FAST sensitivity analysis

Because the primary FAST analyses used absolute tissue volumes, a sensitivity analysis was performed to determine whether the observed group differences were retained after accounting for individual variation in intracranial size. Total intracranial volume values were obtained from the participant metadata accompanying the public source dataset. These values were verified against the EstimatedTotalIntraCranialVol entries in the released FreeSurfer aseg.stats files for all 42 participants and corresponded to FreeSurfer estimated total intracranial volume (eTIV), with the participant-table values rounded to three decimal places. FreeSurfer eTIV is an automated, atlas-scaling-based estimate of intracranial size (Buckner et al., 2004).

Mean eTIV was 1669.0 ± 128.0 $cm^3$ in American football players, 1746.0 ± 105.8 $cm^3$ in cross-country runners, and 1692.4 ± 78.6 $cm^3$ in non-athletes. The three-group Welch analysis did not identify a group difference in eTIV, $F(2, 24.20) = 1.94$, $p = .165$. Across all participants, eTIV was positively associated with each of the four absolute FAST-derived tissue-volume measures, with Pearson correlations ranging from $r = .700$ to .930. These associations indicated substantial covariation between intracranial size and the absolute tissue-volume measurements and motivated examination of the eTIV-adjusted models.

For each FAST outcome, eTIV was converted from $mm^3$ to $cm^3$ and grand-mean centered by subtracting the sample mean of 1703.345 $cm^3$ from each participant's eTIV value. An ordinary least-squares regression model was then fitted with group and centered eTIV as predictors. HC3 heteroskedasticity-consistent covariance estimates were used for inference to retain robustness to unequal residual variances and unequal group sizes (MacKinnon & White, 1985). Homogeneity of regression slopes was evaluated before fitting the common-slope models by including group-by-eTIV interaction terms. No interaction was identified for CSF, $F(2, 36) = 0.18$, $p = .834$; gray matter, $F(2, 36) = 0.42$, $p = .659$; white matter, $F(2, 36) = 0.22$, $p = .804$; or total segmented tissue volume, $F(2, 36) = 1.08$, $p = .349$. Common-slope models were therefore used for the adjusted sensitivity analysis.

The adjusted omnibus group effect was evaluated for each tissue-volume measure using a joint HC3-robust Wald $F$ test. Pairwise adjusted group contrasts were evaluated at the grand mean eTIV of 1703.345 $cm^3$ using robust 95% confidence intervals. Benjamini–Hochberg FDR correction was applied across the four adjusted omnibus tests and separately across the 12 adjusted pairwise contrasts. The adjusted analysis retained the corrected CSF group effect and additionally identified a corrected group effect for white matter, whereas gray matter and total segmented tissue volume did not show corrected adjusted group effects. The detailed adjusted pairwise contrasts are reported in Supplementary Table S5.

**Supplementary Table S5:** eTIV-adjusted group comparisons of FAST-derived tissue volumes. FAST-derived absolute tissue volumes were modeled as a function of group and mean-centered FreeSurfer estimated total intracranial volume (eTIV), with inference based on HC3 heteroskedasticity-consistent covariance estimates. Pairwise adjusted group contrasts were evaluated at the grand mean eTIV (1703.345 $cm^3$). Adjusted mean differences are expressed as the first-named group minus the second-named group in $cm^3$ and are accompanied by robust 95% confidence intervals. Benjamini–Hochberg false discovery rate (FDR) correction was applied across the complete family of 12 adjusted pairwise contrasts. Positive adjusted mean differences indicate greater volume in the first-named group. CI = confidence interval.

| **Tissue-volume measure** | **Contrast** | **Adjusted mean difference ($cm^3$) [95% CI]** | **HC3-robust statistic** | ***p* value** | **FDR *q* value** |
|---|---|---|---|---|---|
| Cerebrospinal fluid volume | American football players – Non-athletes | 20.73 [6.64, 34.82] | $t(38) = 2.98$ | $p = .005$ | $q = .015$ |
| Cerebrospinal fluid volume | Cross-country runners – Non-athletes | 41.52 [24.70, 58.34] | $t(38) = 5.00$ | $p < .001$ | $q < .001$ |
| Cerebrospinal fluid volume | American football players – Cross-country runners | -20.79 [-40.16, -1.43] | $t(38) = -2.17$ | $p = .036$ | $q = .086$ |
| Gray matter volume | American football players – Non-athletes | -6.67 [-40.71, 27.36] | $t(38) = -0.40$ | $p = .694$ | $q = .880$ |
| Gray matter volume | Cross-country runners – Non-athletes | -4.94 [-40.65, 30.76] | $t(38) = -0.28$ | $p = .781$ | $q = .880$ |
| Gray matter volume | American football players – Cross-country runners | -1.73 [-24.72, 21.26] | $t(38) = -0.15$ | $p = .880$ | $q = .880$ |
| White matter volume | American football players – Non-athletes | 1.75 [-14.06, 17.57] | $t(38) = 0.22$ | $p = .824$ | $q = .880$ |
| White matter volume | Cross-country runners – Non-athletes | -27.29 [-40.58, -14.00] | $t(38) = -4.16$ | $p < .001$ | $q = .001$ |
| White matter volume | American football players – Cross-country runners | 29.05 [11.29, 46.80] | $t(38) = 3.31$ | $p = .002$ | $q = .008$ |
| Total segmented tissue volume | American football players – Non-athletes | 15.81 [-15.87, 47.49] | $t(38) = 1.01$ | $p = .319$ | $q = .638$ |
| Total segmented tissue volume | Cross-country runners – Non-athletes | 9.28 [-25.31, 43.88] | $t(38) = 0.54$ | $p = .590$ | $q = .880$ |
| Total segmented tissue volume | American football players – Cross-country runners | 6.53 [-24.61, 37.66] | $t(38) = 0.42$ | $p = .674$ | $q = .880$ |
